# Characterization and pharmacological modulation of Alzheimer’s disease-associated human microglial states

**DOI:** 10.64898/2026.08.26.747247

**Authors:** Gerardo Garcia-Diaz Barriga, Daniel Rosebrock, Henrik Renner, Isabelle Meyer, Georgina Peñalosa-Ruiz, Mariia M Firulyova, Malte Simon, Tim Yang, Giulia Maia Serratto, Fabrizia Zoppetti, Wynona Müller, Anastasia Illarionova, Katharina Heise, Rainer Kuhn, Heinz von der Kammer, Bastian Zimmer, Doris Gruber-Schoffnegger

## Abstract

Microglia are central mediators of Alzheimer’s disease (AD) pathogenesis, yet the mechanisms driving disease-associated microglial states and their therapeutic modulation remain poorly understood. Here, we integrated single-nucleus transcriptomic datasets across the AD spectrum and identified disease- and lipid-associated microglia (DLaM) as a major AD-enriched population linked to genetic risk, neuropathology and cognitive decline. To model this state experimentally, we screened AD-relevant perturbations in human induced pluripotent stem cell (hiPSC)-derived microglia and found that ferric ammonium citrate (FAC) reproducibly induced a DLaM-like state characterized by lipid accumulation, lysosomal dysfunction and impaired Aβ phagocytosis. Using a transcriptomics-based state-reversion screen, we identified LY2090314 as a potent modulator that restored microglial function and induced a distinct lysosomal-metabolic state. These findings establish a framework for transcriptomic disease-state-guided therapeutic discovery in AD.

## Introduction

Alzheimer’s disease (AD) is the leading cause of dementia and is characterized by amyloid-β (Aβ) deposition, tau pathology, neurodegeneration, and progressive cognitive decline. Recent genome-wide association studies (GWAS) have shown that many of the strongest AD risk loci are preferentially expressed in microglial cells establishing them as central contributors to disease pathogenesis^1,2^. Microglial heterogeneity in the human AD brain has been characterized transcriptomically at the single-cell level in multiple landmark studies, identifying disease-associated populations enriched for genes involved in lipid metabolism, lysosomal function, oxidative stress responses and phago-lysosomal remodeling^3–6^. Importantly, these analyses suggest that classical inflammatory activation of microglia alone is insufficient to account for AD pathogenesis^3,6^.

Despite identification of disease-associated microglial populations, translating human transcriptomic observations into experimentally tractable models has remained a major challenge. To address this, human induced pluripotent stem cell (hiPSC)-derived microglia have emerged as powerful experimental *in vitro* system to study AD biology. However, current *in vitro* microglia models rely on inflammatory stimulation, epigenetic perturbations, or transcription factor overexpression^3,7–9^ that incompletely reproduce molecular programs observed in patient tissue and, in addition, face scalability limitations hindering their use for therapeutic discovery. This highlights the need for strategies that directly connect human disease signatures to hiPSC microglial models. Recent advances in perturbational genomics and transcriptomic drug discovery leverage high-dimensional gene-expression signatures rather than individual molecular markers to define cellular phenotypes^10–13^. In this framework, disease-associated transcriptional programs can serve as quantitative phenotypes for identifying compounds that induce or reverse pathological cell states. Such approaches have proven effective across multiple biological systems^14,15^, yet their application to human neurodegenerative disease-associated microglial states has remained limited^16^.

Here, we address the issue by combining large-scale human single-nucleus transcriptomics with perturbational modeling and transcriptomic-reversal screening in hiPSC microglia. Using newly generated and published human brain single-nuclei transcriptomics datasets, we identified disease- and lipid-associated microglia (DLaM) as a major AD-associated population linked to disease progression. Importantly, we produced a human iPSC-derived microglia model in which iron-induced metabolic stress reproducibly generated a DLaM-like state closely correlating with AD patient signatures, displaying lipid droplet accumulation, impaired lysosomal degradation and reduced Aβ phagocytosis. Finally, using this model, we developed a transcriptomics-based reversal screen evaluating compounds with known mechanisms of action. Notably, the small molecule LY2090314, a previously described glycogen synthase kinase 3 (GSK3) inhibitor^17^, recovered microglial function while inducing a distinct lysosomal-metabolic transcriptional program. Together, these findings establish a framework for translating human disease-associated cellular states into tractable screening paradigms and demonstrate how transcriptomic state reversal can uncover therapeutic mechanisms for reprogramming maladaptive microglial responses in Alzheimer’s disease.

## Results

### Construction of a large-scale single-nucleus transcriptomic atlas of Alzheimer’s disease

To define cell type–specific molecular alterations in Alzheimer’s disease (AD), we generated a single-nucleus RNA sequencing (snRNA-seq) atlas from postmortem frontal and temporal cortex of AD patients (n = 30) and age-matched controls (n = 42). Samples were selected to represent a balanced distribution of neuropathological and clinical diagnoses (Supplementary Table 1). To increase statistical power and capture disease heterogeneity, we integrated these data with previously published snRNA-seq datasets (Fig. 1a; Methods). Following metadata harmonization, individuals were assigned to joint diagnostic categories (CTR, MCI−, MCI+, AD and resilient) based on concordance between neuropathological and clinical criteria (Table 1).

**Table 1:**

|  | Clinical | Neuropathological |
| --- | --- | --- |
| Scores and Criteria | Clinical Dementia Rating (CDR) | Tau: BraakStage |
|  | CTR: MCI: AD: | Amyloid: Thal, CERAD |
|  | 0 0.5-1 >1 | combined: ABC score |
|  | cogdx (ROSMAP) | CTR: AD: |
|  | CTR: MCI: AD: | "Not", "Intermediate", |
|  | 1 <2≤3 ≥4 | "Low" "High" |
| Joint_Diagnosis | Clinical | Neuropathological |
| CTR | CTR | CTR |
| MCI- | MCI | CTR |
| MCI+ | MCI | AD |
| resilient | CTR | AD |
| AD | AD | AD |

**Fig. 1.**
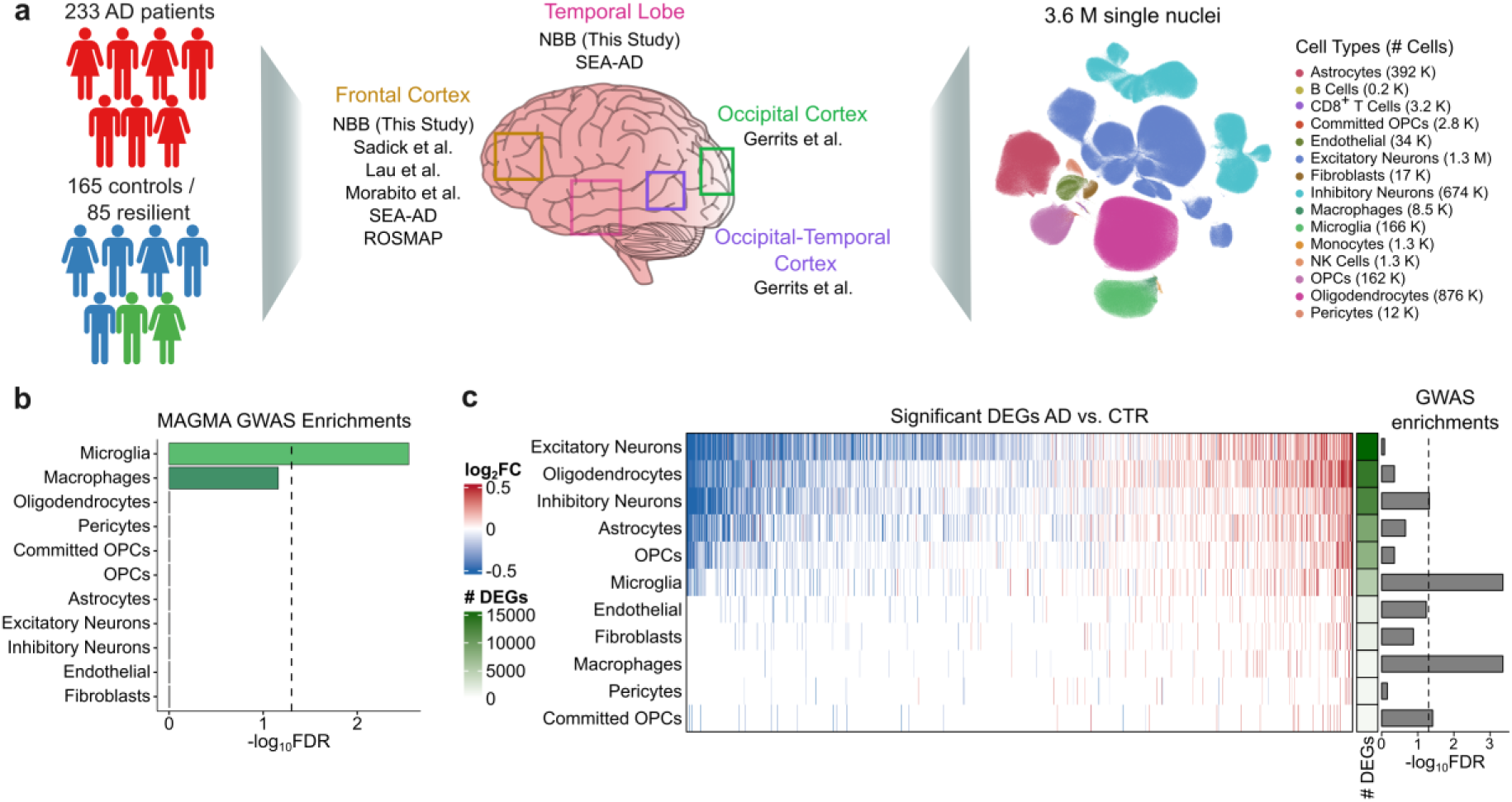
Large-scale single-nucleus transcriptomic atlas of Alzheimer’s disease across human cortex. **a)** Study design and data integration with overview of patient cohort (left), available brain regions and data sources (middle), and UMAP highlighting major cell types identified by reference-based annotation (right). **b)** Enrichment of genetic risk across all cell types. Multi-marker analysis of genomic annotation (MAGMA) applied to Alzheimer’s disease genome-wide association study (GWAS) data, using the top 10% of marker genes per cell type (FDR < 0.05). c) Heatmap showing AD vs. CTR DEGs within each cell type (as determined by limma-voom pipeline, see Methods). The barplot on the right displays results from overrepresentation of cell type-wise DEGs against Bellenguez et. al. GWAS significant genes (hypergeometric test with Benjamini-Hochberg correction, dashed line at FDR = 0.05).

The combined dataset comprised 233 AD cases, 165 cognitively normal controls and 85 cognitively resilient individuals resulting in a dataset of 3.63 million high-quality nuclei from frontal, temporal, occipital and occipito-temporal cortices. Cohort characteristics were consistent with established AD epidemiology^18^ (Extended Data Fig. 1a). Reference-based annotation^19^ identified 15 major cell types (Fig. 1a). Cell-type identities were supported by canonical marker expression (Extended Data Fig. 1b; Supplementary Table 2) being restricted to expected cell classes. Corresponding cell types integrated well across individual datasets and brain regions (Extended Data Fig. 1c). Projection of CTR and AD samples onto a shared embedding revealed preserved global organization but localized differences in cell density (Extended Data Fig. 1d), visible within both neuronal and glial populations.

### Disease- and lipid-associated microglia are enriched in Alzheimer’s disease

To assess genetic relevance of individual cell types, we applied multi-marker analysis of genomic annotation (MAGMA) using the top 10% of marker genes per cell type and AD GWAS summary statistics^1^ (Fig. 1b; Supplementary Table 3). Based on MAGMA, microglia showed the strongest enrichment for AD risk. This was further supported by GWAS hits being most significantly enriched among differentially expressed genes (DEGs) comparing AD against CTR samples within microglia and macrophages (Fig. 1c; Supplementary Tables 4/5), despite these cell types having fewer DEGs relative to other cell types.

To resolve microglial heterogeneity and relate identified states to neurodegeneration-associated microglial phenotypes ^20^, we performed clustering of 166,321 microglial identifying 19 clusters (Fig. 2a, Proliferating cluster (Prolif_17) was excluded, see Extended Data Fig. 2a). Clusters were annotated based on canonical markers^6^ and were evenly distributed across studies, regions, and sex, indicating limited confounding effects (Extended Data Fig. 2b, Supplementary Table 6). The transcriptional landscape formed a continuum from homeostatic to disease- and stress-associated states. Surveilling clusters corresponding to homeostatic microglial states were defined by expression of *CX3CR1*, *P2RY12* and *PRDM1* (Fig. 2b). Pathway enrichment indicated signaling via P2Y receptors, GPCRs and TYROBP-associated networks (Fig. 2c, Supplementary Table 7). Transitional clusters (Surveilling-Enhanced Redox 0 and Surveilling-Disease/Lipid-Associated 13) co-expressed homeostatic as well as metabolic- and stress-related genes, including ribosomal and redox-associated transcripts, suggesting intermediate states rather than discrete identities. Disease- and lipid-associated microglia (DLaM; clusters 8 and 15) were defined by expression of *APOE*, *TREM2*, *LPL*, *GPNMB,* and *PTPRG*, consistent with previously described Alzheimer’s disease-associated microglia states^3,6^. While both DLaM8 and DLaM15 exhibited a selective enrichment for lipid-associated pathways, DLaM8 exhibited a specific up-regulation of receptor-mediated endocytosis, while DLaM15 specifically increased eosinophil migration (Fig. 2c). Inflammatory clusters (1, 6 and 7) expressed cytokine-related genes (*IL1B*, *IL21R*, *HSPA5*) and overlapped with markers reported for Human Alzheimer’s Microglia (HAM)^21^. Similarly, reactive cluster 10 was also characterized by upregulation of genes related to inflammatory responses (*CXCR4/C5AR1*) and nuclear receptors (*NR4A3*). Non-canonical inflammatory and stress-associated clusters (12 – *HSP90AA1/FKBP4*, 14 and 16 – *SERPINE1/GPR183*) occupied central positions within the UMAP, suggesting convergence of multiple stress-related states.

**Fig. 2.**
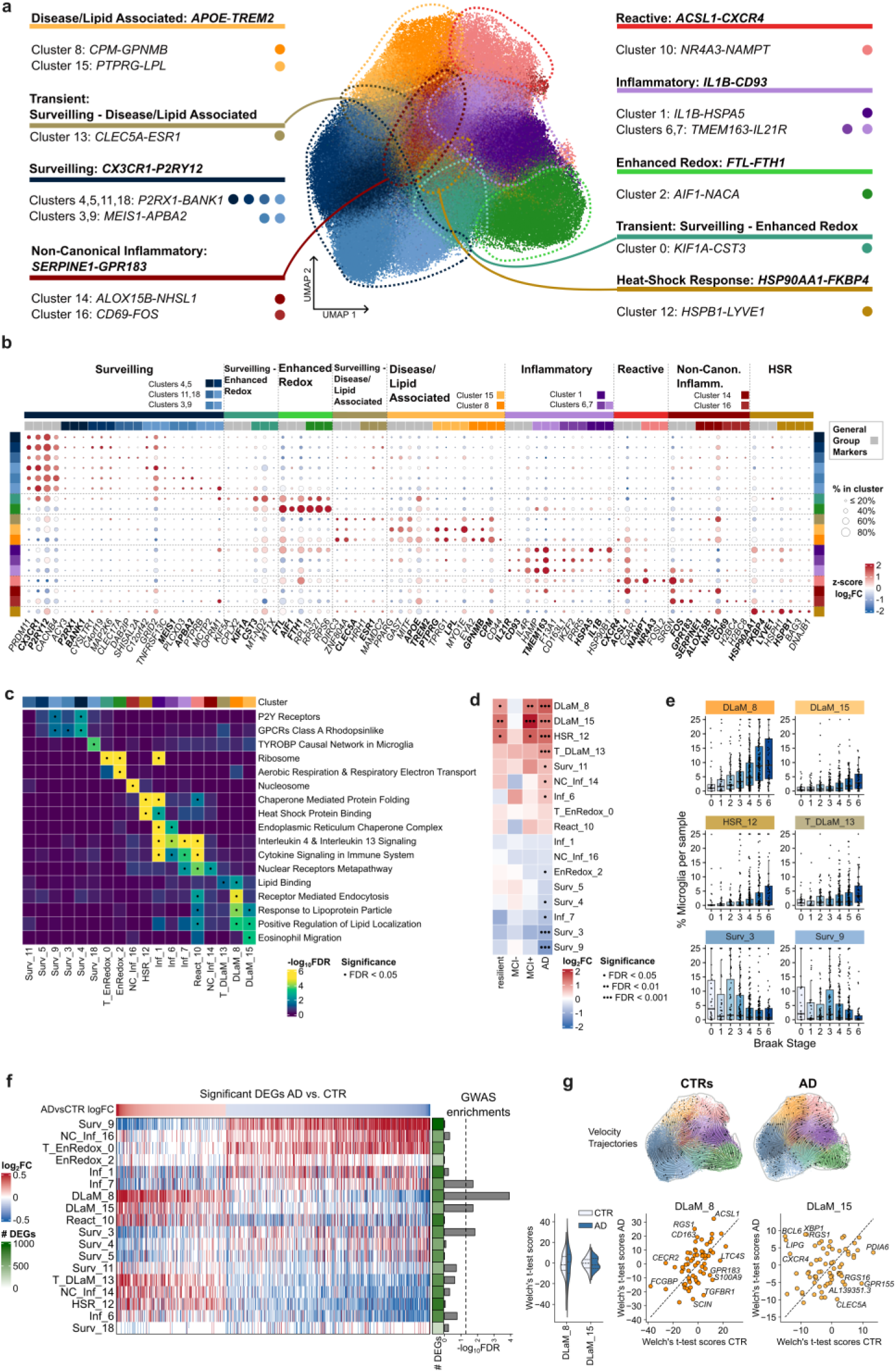
Disease- and lipid-associated microglia are enriched in Alzheimer’s disease. **a)** UMAP based on 166,000 microglia nuclei showing 19 clusters identified using Leiden clustering and resolution 1.2 (a small proliferative cluster 17 was excluded; see Extended Data Fig. 2a) with key markers annotated for each cluster. **b)** Dotplot graph showing key microglia subpopulation markers and their relative expression across microglia states (normalized log2fold change (FC) of cluster vs. all others), as well as percentage of marker expressing cells in cluster (size). Grey sidebar coloring indicates main microglia state markers; individual cluster markers are highlighted by additional colors. **c)** Pathway enrichment differentiating microglia clusters. Heatmap highlighting pathways significantly enriched in different microglia subpopulations (hypergeometric test with Benjamini-Hochberg correction, (·) = FDR < 0.05). **d)** Associations with joint neuropathology/clinical diagnosis (empirical Bayes quasi-likelihood F-tests corrected for confounders, (·) = FDR < 0.05, (··) = FDR < 0.01, (···) = FDR < 0.001). **e)** Box plots show correlation of enrichment/depletion of the respective clusters with increasing Braak stage. **f)** Heatmap of the overlap of microglia cluster markers with AD vs CTR microglial DEGs. Microglia cluster markers genes (FDR < 0.05, each vs other clusters) were filtered for AD vs. CTR DEGs within microglia (determined by limma-voom pipeline FDR < 0.05, see Methods) sorted by AD vs. CTR log2FC estimates, color scale represents cluster comparisons log2FC. Bar plots (right) show enrichment of filtered cluster DEGs against Bellenguez et al. AD GWAS set (hypergeometric test with Benjamini–Hochberg multiple hypothesis correction, dashed line: FDR = 0.05). **g)** snRNA-seq velocity and trajectory analysis. Visualizations on UMAP for CTR and AD depict velocity embeddings through microglial states (top). Quantification of velocity enrichments by gene as estimated by Welch’s t-test comparing DLaM8 and DLaM15 clusters, respectively, against all other clusters (bottom left), scatterplot of Welch’s t-test scores by gene comparing AD vs CTR within the DLaM8 cluster (bottom center) and DLaM15 cluster (bottom right). Genes with highest deviation from the diagonal (dashed line) are labeled.

The cell density landscape of microglia within CTR and AD donors displayed a marked shift in microglia composition within disease (Extended Data Fig. 2c). Concordantly, cluster abundance analysis revealed significant enrichment of DLaM8, DLaM15 and HSR12 in AD donors and with increasing Braak stage (Fig. 2d/e; Extended Data Fig. 2d and Supplementary Table 8). In contrast, Surv3 and Surv9 were reduced with increasing Braak stages and were overall significantly depleted in AD. These analyses consistently identified DLaM microglia as most closely linked to AD pathology and cognitive impairment. Comparison with published datasets (Extended Data Fig. 2e/f) showed concordance between identified clusters and previously reported microglial states^3,6^. DLaM8 aligned with lipid-associated states linked to amyloid pathology (Mic12 in Green et al.), whereas DLaM15 corresponded to tau-associated states linked to cognitive decline (Mic13 in Green et al.). Enrichment analysis of AD GWAS hits among cluster-specific DEGs showed strongest enrichment in DLaM8, with additional significant enrichment in DLaM15, Inf7 and Surv3 clusters (Fig. 2f, Supplementary Tables 5/6). RNA velocity overlaid on CTR and AD UMAPs (Fig. 2g) revealed distinct microglial dynamics. In CTR, trajectories showed a steady flow from Enhanced Redox towards either Surveilling or Inflammatory/Reactive states. In AD, flows were fragmented, with a transition from Inflammatory clusters to the transient Surveilling–Enhanced Redox cluster and an increased flow from Reactive to DLaM microglia. We next compared relative RNA velocity enrichments of DLaM 8 and 15 clusters within AD and CTR samples. DLaM15 microglia exhibited a smaller spread of RNA velocity enrichments, indicating a weaker directional transcriptional activity more reminiscent of a terminal state. Finally, AD samples exhibited increased RNA velocity enrichments of *LIPG* and *ACSL1*, both involved in lipid transport and metabolism, further implicating these processes in disease.

### Ferric ammonium citrate (FAC) induces a disease- and lipid-associated state in hiPSC-derived microglia

As DLaM15 microglia appeared to represent a detrimental end-stage phenotype, we sought to recapitulate this state *in vitro* to enable the identification of mechanisms regulating its maintenance. To model disease-associated microglial states, we exposed human iPSC-derived microglia to perturbations mimicking AD-relevant processes, including oxidative stress, metabolic dysfunction, inflammation and amyloid exposure (Fig. 3a). Transcriptomic profiling using 384-well plate-based RNA sequencing (*ScreenSeq*)^22^ (for details, see Methods) revealed a clear separation of stimuli in principal component space (Fig. 3b). Brain cell-type-specific gene expression analysis (BRETIGEA) deconvolution^23^ using microglia cluster markers (Supplementary Table 6, see Methods), identified FAC as the only stimulus inducing a DLaM15 transcriptional profile, whereas metabolic perturbations preferentially induced a DLaM8 profile (Fig. 3c; Extended Data Fig. 3a). We observed consistent induction of microglia disease-associated states by deconvolving the same data based on microglia clusters previously described^3,6^ (Extended Data Fig. 3b/c). DEGs induced by FAC stimulation overlapped with key DLaM markers, including *LPL* and *PPARG*, key modulators of lipid metabolism, the well-known AD risk genes *TREM2* and *APOE*, as well as *SPP1 and CD9*, one of the signature genes used to identify DAM^24^ (Fig. 3d, Supplementary Table 9). Pathway analysis (Fig. 3e, Supplementary Table 10) showed enrichment of lipid binding and lipoprotein response pathways including those containing AD risk genes (Extended Data Fig. 3d). Other stimuli, including LPS/IFNγ or Aβ aggregates, induced inflammatory states (Fig. 3c) characterized by cytokine and interferon signaling (Fig. 3e, Extended Data Fig. 3e/f). Most perturbations also showed downregulation of cell cycle and ribosomal pathways (Fig. 3e). Comparison with published *in vitro* disease models (Fig. 3f, Extended Data Fig. 4) demonstrated that the FAC-induced transcriptional signature showed the strongest concordance with DLaM15 and other published AD-associated microglia states compared to models based on MITF overexpression, HDAC inhibition or *BHLHE40/41* transcription factor knockout^7–9^. These results further strengthen the finding that FAC stimulation mimics a disease-associated state in hiPSC microglia with higher specificity of FAC stimulation to induce this population compared to other state-of-the-art models.

**Fig. 3.**
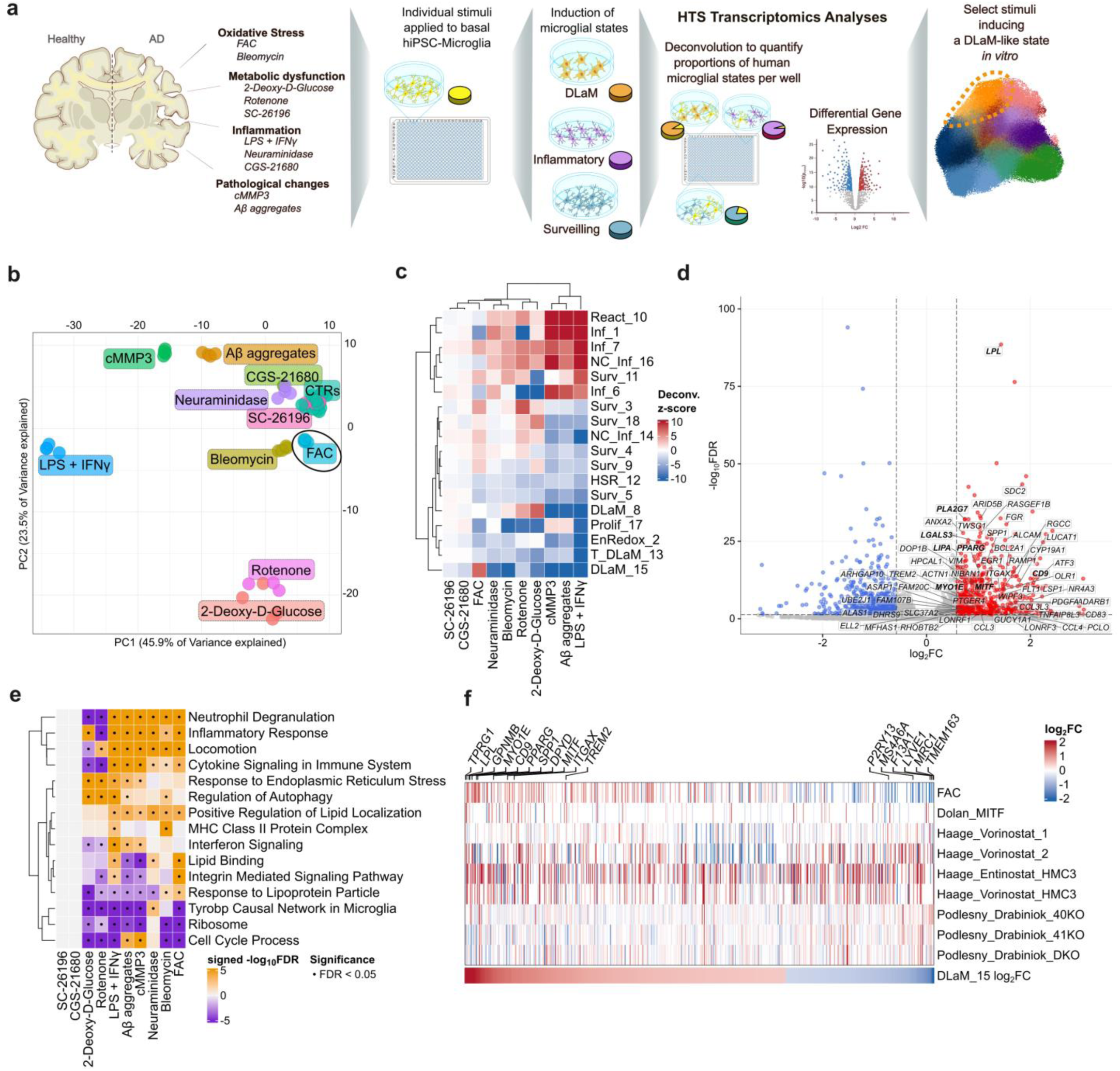
Modeling disease- and lipid-associated states in human iPSC-derived microglia. **a)** Experimental pipeline. Schematic of the workflow used to profile the impact of different disease-relevant stimuli on the transcriptomic state of hiPSC-derived microglia and correlating them to patient data. b) Global transcriptional variation visualized by PCA of the top 2,000 variable genes. c) Deconvolution to human microglial states. Heatmap of BRETIGEA based normalized z-scores (see Methods) using microglia cluster markers from Fig. 2. d) FAC-induced transcriptional signature shown as Volcano plot FAC vs. solvent control (as determined by limma-voom pipeline, see Methods), labeled genes are markers for DLaM 15 e) Pathway enrichment across stimuli. The color of each cell represents median log2FC across all DEGs of each stimulation vs. CTR (as determined by limma-voom, FDR < 0.05, no log2FC filtration) of a given pathway, significance derived from hypergeometric test with Benjamini– Hochberg correction of overrepresentation of DEGs in pathway sets (·) = FDR < 0.05. f) Heatmap showing overlap of patient DLaM15 cluster markers with log2FC estimates of FAC stimulated hiPSC microglia and other published perturbations (MITF overexpression; HDAC inhibitor treatments; BHLHE40/41 single and double knockouts). Genes are grouped by DLaM15 cluster marker FC (bottom, FDR <0.05 |log2FC| > 0.5). FAC DEGs (FDR < 0.05 from limma-voom pipeline t-statistic test with BH correction), other system DEGs as reported in respective references7-9.

### FAC-treated hiPSC-derived microglia contain a higher percentage of a “DLaM15-like” population and mimic AD microglial dysfunction

Single-cell RNA-seq of FAC-treated and untreated microglia identified three major populations: control-like, cycling, and DLaM15-like, as well as two small clusters, one characterized by IFN-signaling (IFN-high) and the other by high expression of chemokines (CCL-high) (Fig. 4a, small non-microglial clusters excluded, see Extended Data Fig. 5a-c). The DLaM15-like cluster was defined by expression of DLaM15 markers *CD9*, *LPL*, *PPARG*, *TREM2* and *TPRG1* (Fig. 4b; Supplementary Table 11). Microglia of the Control cluster expressed markers of surveilling populations (*P2RY6, IGF1, DLEU7*), as well as genes expressed in other microglia clusters (*LILRB2* – React10, *MS4A6A* – DLaM8). The expression level of Control cluster markers was lower in FAC-treated microglia in comparison to control microglia (Fig. 4b), suggesting that the surveilling capabilities of DLaM15-like microglia might be reduced. FAC treatment increased the proportion of DLaM15-like microglia from 32% to 65% and reduced Control microglia from 53% to 27% (Fig. 4c). Module scoring using human microglia cluster markers confirmed higher DLaM15 module scores in FAC-treated DLaM15-like microglia and higher Surv5 module scores in a subset of Control microglia (Fig. 4d).

**Fig. 4.**
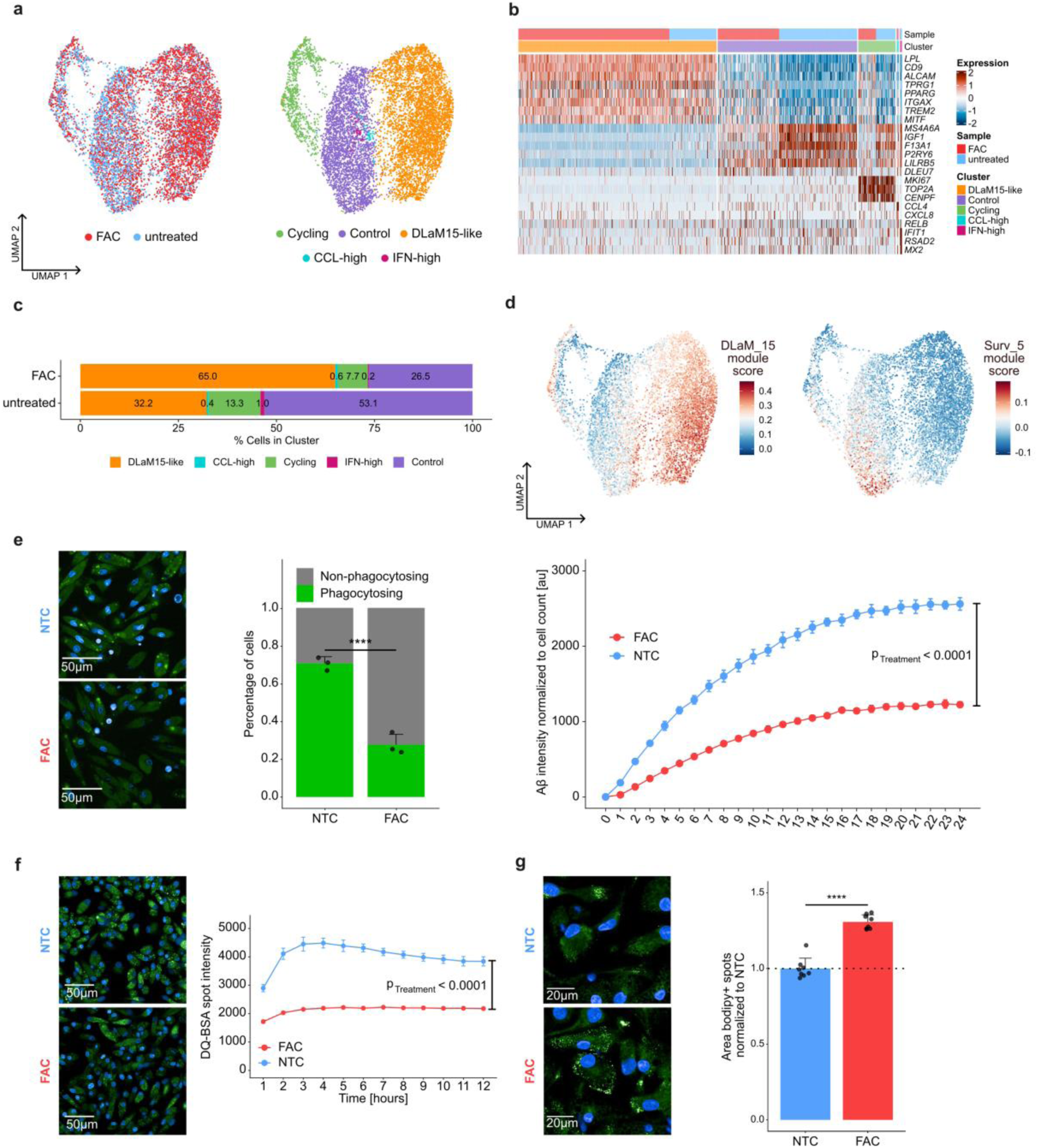
FAC-treated human iPSC microglia recapitulate Alzheimer’s disease–like microglial dysfunction. **a)** Single-cell RNA sequencing of hiPSC microglia. UMAPs of untreated and FAC-treated hiPSC microglia (left: colored by sample; right: by clusters). Mesenchymal and mast cells, which constituted <1% of the cells were excluded from the UMAP (see Extended Data Fig. 5a). **b)** Heatmap of cluster markers and their expression in individual cells grouped by cluster and sample. **c)** Stacked bars visualizing compositional changes of hiPSC microglia cultures with FAC. **d)** UMAP overlays of gene-set module scores for DLaM15 and Surveilling 5 (Surv5). **e)** Aβ phagocytosis. Left: representative images (green: pHrodo-Aβ, blue: nuclei). Middle: Stacked bars showing percentage of phagocytosing and non-phagocytosing cells per condition (n = 3; Phagocytosing % comparison, two-tailed Welch’s unpaired t-test, *** = p <0.001). Right: 24-hour time course of Aβ-intensity normalized to cell count (n = 4, two-way RM ANOVA with Geisser-Greenhouse’s correction, main Timepoint effect F[2.932,17.59] = 3115, p <0.0001, η2p= 0.562; main Treatment effect F[1,6] = 633.4, p <0.0001, η2p = 0.374 and interaction Timepoint × Treatment F[2.932,17.59] = 326.9, p <0.0001, η2p= 0.059). **f)** Lysosomal activity. Left: representative images (green: DQ-BSA, blue: nuclei). Right: DQ-BSA pulse-chase (uptake titrated to match between conditions; see Extended Data Fig. 5d), (n = 3; two-way RM ANOVA with Geisser-Greenhouse’s correction, main Timepoint effect F[1.999,7.995] = 41.56, p <0.0001, η2p= 0.067; main Treatment effect F[1,4] =476.1, p <0.0001, η2p= 0.894 and interaction Timepoint × Treatment F[1.999,7.995] = 14.57, p =0.0022, η2p= 0.023.) g) Lipid droplets. Left: representative images (green: BODIPY, blue: nuclei). Right: BODIPY staining normalized to untreated controls (NTC), (n=8, **** = p < 0.0001 using a two-tailed Welch’s unpaired t-test). All data are presented as mean ± SD.

Functionally, FAC-treated microglia exhibited reduced Aβ phagocytosis over a 24 h time-course, with the proportion of phagocytic cells decreasing from 71% to 28% at the 6 h-timepoint (Fig. 4e). Lysosomal degradation, assessed by DQ-BSA pulse-chase assay, was reduced in FAC-treated cells (Fig. 4f and Extended Data Fig. 5d) while lipid droplet abundance increased by 30% (Fig. 4g). These functional changes were consistent with the transcriptional profile of DLaM-like microglia, including enrichment of lipid-related pathways and reduced expression of genes associated with surveilling function.

### Pharmacological modulation of FAC-treated hiPSC-derived microglia produces a diverse set of transcriptional states diverging from DLaM15

We confirmed the robustness of the DLaM15-like transcriptomic state by measuring reproducibility of the most consistently up- and downregulated FAC-induced DEGs over seven independent experiments (Extended Data Fig. 6). To assess whether DLaM15-like hiPSC microglia could be modulated pharmacologically, we performed a compound screen in a reversion paradigm (Fig. 5a, see Methods), applying 54 compounds targeting over 15 distinct mechanisms (Fig. 5a-b, Supplementary Table 12). UMAP visualization showed that most compounds had no effect on the transcriptome and clustered with FAC-treated samples, while subsets induced distinct transcriptional shifts (Fig. 5b, Extended Data Fig. 7a). A signature reversal score (σ, see Methods) was used to quantify the extent to which compounds reverted the FAC-induced transcriptional program (Fig. 5c). SYK inhibitors showed the strongest reversal, with a concentration-dependent increase in the number of DEGs elicited that corresponded well with the reported cellular IC_50_ (Fig. 5d). BTK inhibitors produced similar effects. Additional partial reversion was observed for PAK inhibitors, TLR agonists and selected GSK3 inhibitors. All remaining compounds showed minimal or no significant effect on the FAC-stimulated microglial signature (Extended Data Fig. 8a). All DEGs for every compound at every tested concentration are summarized in Supplementary Table 13.

**Fig. 5.**
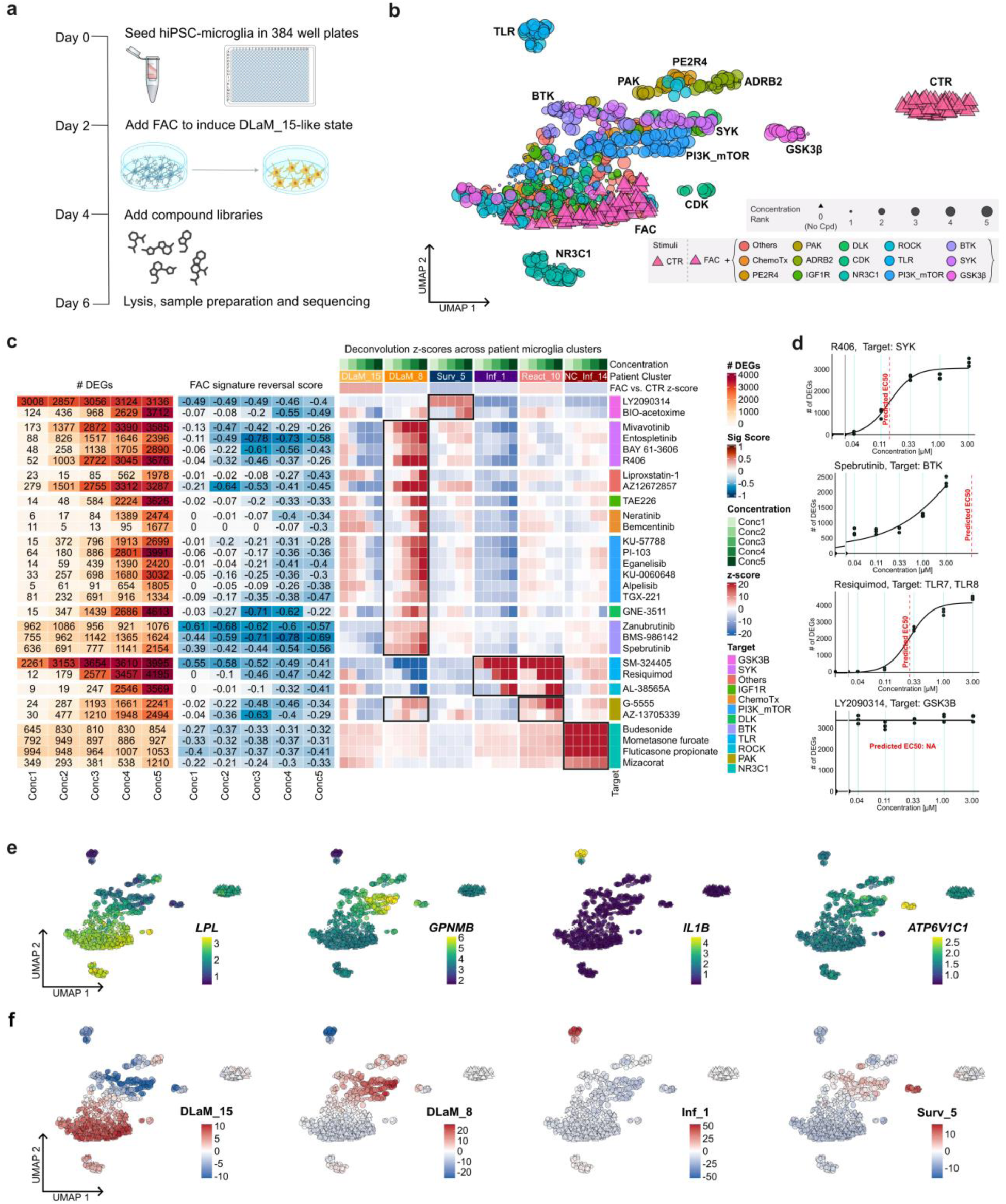
Transcriptomics-based small molecule screening identifies pharmacological modulators of DLaM-like human iPSC microglia. **a)** Screen design. Timeline and workflow of the transcriptional state modulator screen. Compound screening was performed using automated pipetting workflows in 384-well plates and in a reversion paradigm. Samples were analyzed via high throughput plate-based transcriptomics. **b)** Global compound effects. UMAP of all compound-induced profiles. **c)** Overall compound effects, signature reversal and compound-induced shifts in hiPSC microglia transcriptional profiles analyzed by deconvolution to patient microglia clusters. Heatmaps summarize number of DEGs (as determined by limma-voom, FDR < 0.05, no log2FC filtration), FAC signature-reversal score (σ), and averaged deconvolution scores across a subset of patient microglia clusters per compound and dose. Compounds were included if they had at least 500 DEGs in at least one concentration and a FAC signature-reversal score of less than −0.3 in at least one concentration. Black boxes indicate patient clusters with anti-correlated deconvolution estimates compared against DLaM_15 that are common to all compounds with the same target. Accompanying annotation bars indicate compound concentrations, patient microglia clusters, and averaged FAC deconvolution z-scores by patient microglia cluster (all top), and annotated compound targets (right). **d)** Dose response transcriptomic changes. Example dose response plots relate the number of DEGs to concentration for selected compounds (rep-free DEG numbers fitted to an exp5 model with non-linear squares regression (see Methods)). **e)** Gene level compound effects. UMAP overlays of selected genes regulated by compounds. **f)** Deconvolution of hiPSC microglia using patient microglia clusters. BRETIGEA-based deconvolution z-scores (see Methods), using microglial cluster markers (as in Fig. 2a) projected onto the experiment’s UMAP.

To relate these results to human microglia states, we applied microglia cluster marker deconvolution as previously. Expectedly, the expression of DLaM15 markers was most pronounced in areas of the UMAP with the highest DLaM15 deconvolution estimates (Fig. 5 c/e/f, Extended Data Fig. 7a-b) which contained FAC-treated samples without the application of compounds. Conversely, the expression of DLaM8 markers and DLaM8 deconvolution estimates were highest in UMAP areas containing FAC-stimulated samples which had been treated with BTK, PI3K_mTOR or SYK inhibitors. Interestingly, almost all compounds that reduced DLaM15 deconvolution values did so at the expense of a marked increase in DLaM8 deconvolution values, with the major exceptions being the GSK3 inhibitor LY2090314, TLR activators and the ROCK inhibitor AL-38565A (Fig. 5c, Extended Data Fig. 8a). The deconvolution estimates for the Inflammatory 1 population, along with the expression of *IL1B*, a main marker of this population, were strongly increased in FAC-stimulated samples treated with TLR activators and AL-38565A, which also clustered in separate “islands” on the UMAP. In contrast, a subset of GSK3 inhibitors including LY2090314 exhibited a strong increase in deconvolution scores for the Surveilling 5 population and were characterized by the specific up-regulation of *ATP6V1C1*, a component of the vacuolar ATPase, in line with the specific up-regulation of vacuolar acidification and endolysosomal pathways in LY2090314- and BIO-acetoxime-treated samples (Extended Data Fig. 8a). More generally, pathway analysis (Extended Data Fig. 8a, Supplementary Table 14) revealed modulation of inflammatory, lipid metabolic and cytoskeletal pathways across compound classes, with varying degrees of specificity. Taken together, these results demonstrate the capability of compounds to modulate an AD microglia state and the powerful utility of transcriptomics as a readout to measure the enhancement or reduction of relevant microglial states derived from patient data.

### Pharmacological modulation of FAC-treated hiPSC-derived microglia with selected hits restores phagocytic function and alters lipid and metabolic pathways

Functional evaluation of compounds modulating the transcriptomic signature of DLaM15-like microglia revealed divergent effects on microglial activity. While TLR agonists increased Aβ phagocytosis (Fig. 6a), SYK and BTK inhibition reduced it (Fig. 6b/c). Treatment with the GSK3 inhibitor LY2090314 increased phagocytosis, reduced lipid droplet accumulation and restored lysosomal degradation in a concentration-dependent manner (Fig. 6d-f). As the screening experiment results suggested saturation of LY2090314’s effects already at the lowest tested concentration (Fig. 5d), we repeated evaluation of transcriptional changes elicited by LY2090314 on FAC-treated hiPSC microglia expanding the concentration range from 0.02 nM up to 10 µM. LY2090314 dose-dependently modulated 1236 genes (Supplementary Table 15, see Methods), including downregulating inflammatory and lipid-associated genes (*LPL*, *PPARG*, *MITF*) and upregulating genes associated with iron export (*SLC40A1*), anti-inflammatory signaling (*IL10*), and lysosomal pathways (*ATP6V0D2*) (Fig. 7a, Extended Data Fig. 9). Pathway analysis confirmed downregulation of inflammatory pathways and upregulation of lysosomal and metabolic processes (Fig. 7b, Supplementary Table 16) at relevant concentrations relative to the published potency of LY2090314^17^. Finally, integration of significantly enriched pathways into a network-based representation revealed coordinated modulation of interconnected biological processes following treatment with LY2090314 (Fig. 7c). Central nodes included oxidative phosphorylation, ROS and RNS production in phagocytes, cytokine signaling in the immune system, ferroptosis, and the TYROBP causal network, further supporting the functional effects LY2090314 treatment had in hiPSC microglia.

**Fig. 6.**
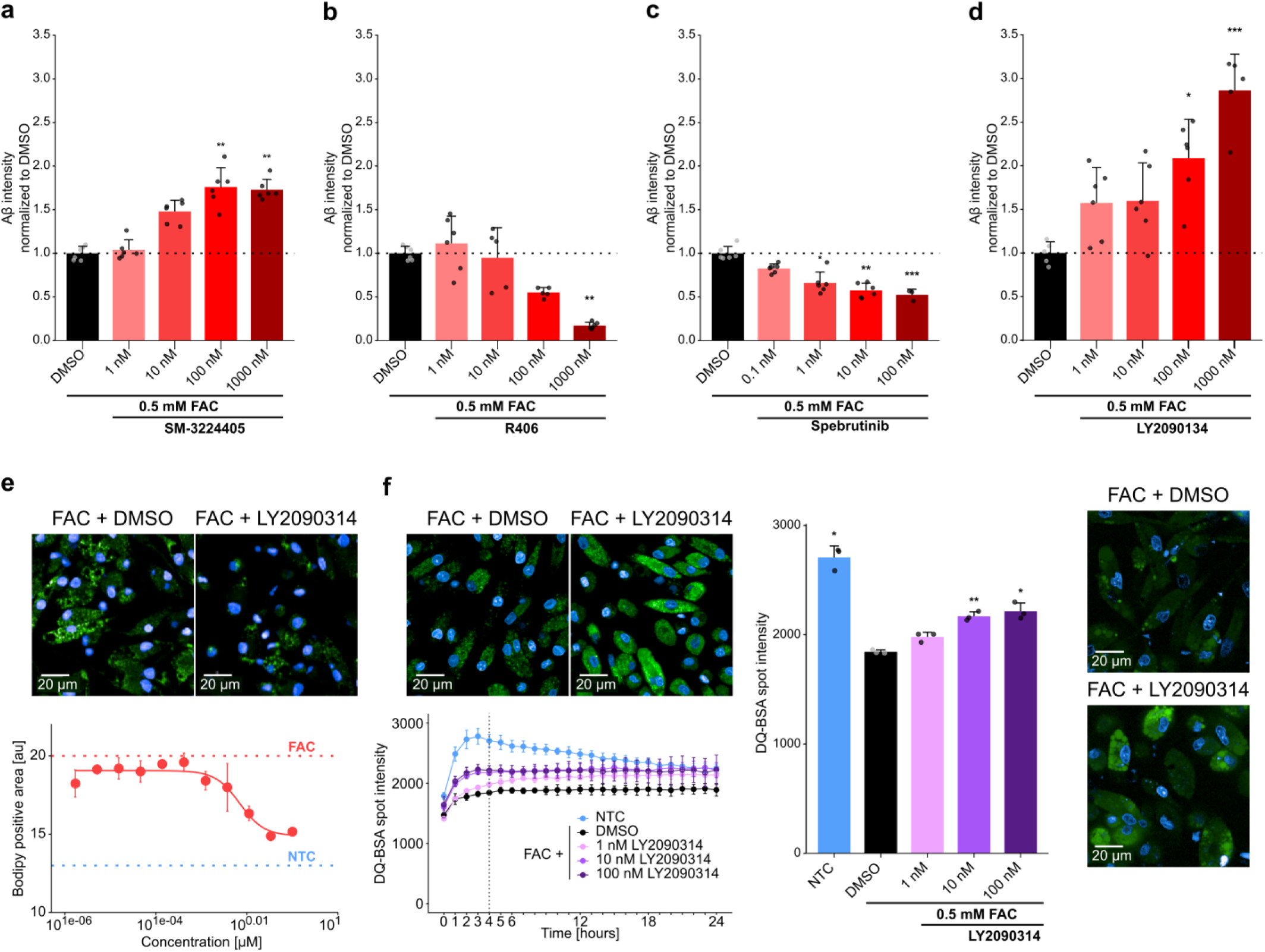
Pharmacological modulation of DLaM-like human iPSC microglia alters cell function. **a - d)** Effects of pharmacological modulation on hiPSC microglia Aβ phagocytosis. Bar plots show Aβ intensity normalized to DMSO controls at the 6-hour timepoint for a) TLR agonist SM-324405, b) SYK inhibitor R406, c) BTK inhibitor spebrutinib, d) GSK3B inhibitor LY2090314. (n ≥ 5 for all compounds, *=p < 0.05, **=p < 0.01 ***=p < 0.001, Kruskal-Wallis test with corrected post-hoc Dunn’s multiple comparison test vs DSMO). Representative example images (bottom of (d)) show pHrodo-Aβ in DMSO controls and 100 nM LY2090314 (blue: nuclei; green: pHrodo-Aβ) **e - f)** LY2090314 effects in hiPSC microglia functional assays. Top: representative example images from the different functional assays; shown are DMSO controls and 100 nM LY2090314 (blue: nuclei; green: BODIPY (e), or DQ-BSA (f)). e) BODIPY quantification concentration response curve for LY2090314 (n = 3), dotted lines indicate DMSO (red) and non-stimulated (without FAC, blue, NTC) levels (data were fitted with a four-parameter log-logistic model. Estimated EC50 = 6.88 × 10⁻³ (95% CI, 0.002 to 0.011), Hill coefficient = 19.1 (95% CI, 18.6 to 19.5), lower asymptote = 1.63, upper asymptote = 14.89). f) DQ-BSA pulse-chase with full time course (left) and detailed comparison at the 4-hour timepoint (right), (n = 3, for time-course: mixed-effects model (REML) with time as a repeated measure and Geisser-Greenhouse’s correction, main Timepoint effect F[1.94,19.37] = 103.5, p <0.0001; main Treatment effect F[4,10] = 17.71, p =0.0002 and interaction Timepoint × Treatment F[7.75,19.37] = 12.29, p <0.0001; post-hoc Tukey’s multiple comparison test vs DMSO is shown for the 4-hours timepoint with *p < 0.05, **p < 0.01). All data are presented as mean ± SD

**Fig. 7.**
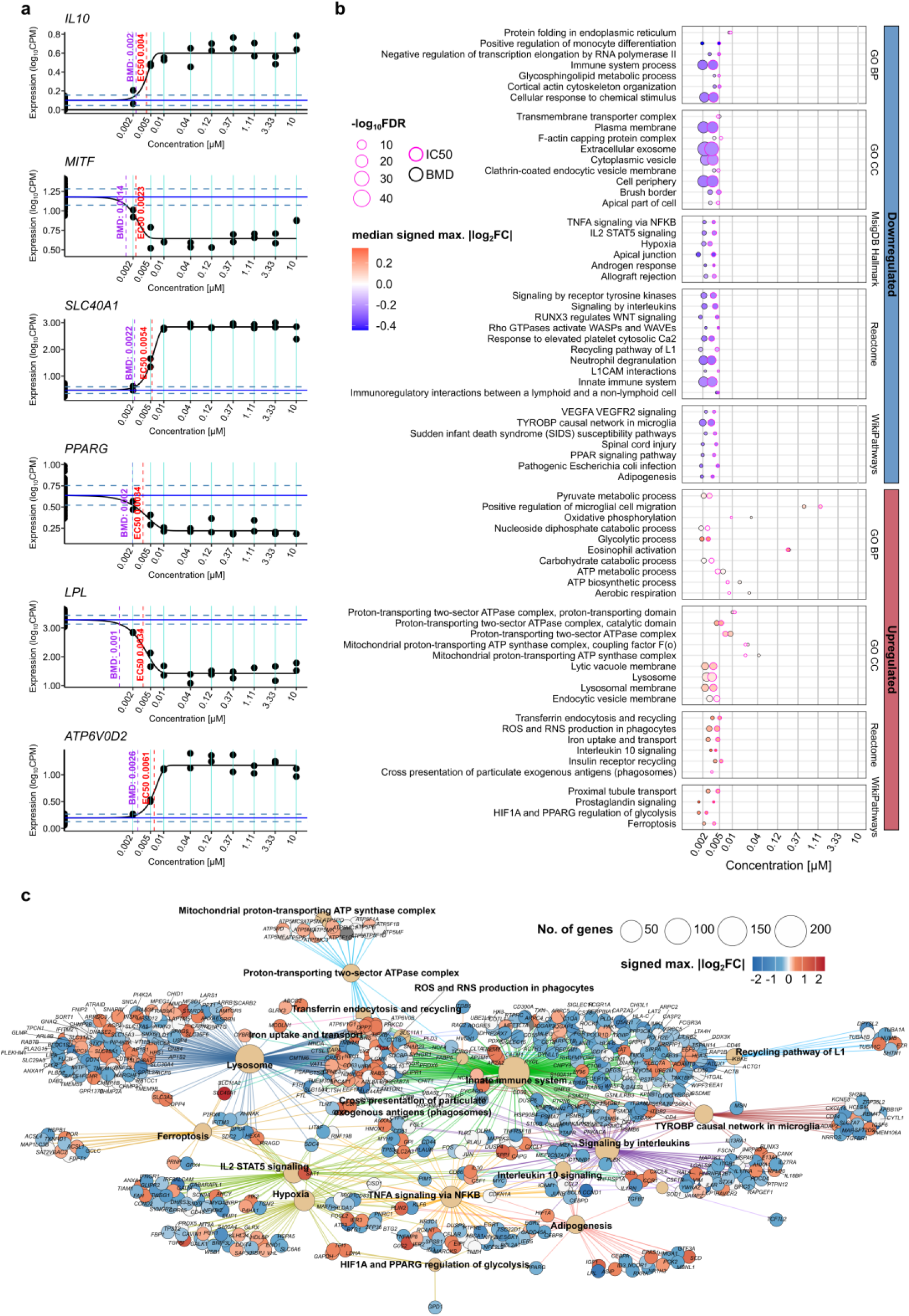
LY20G0314 remodels the transcriptomic landscape of DLaM-like human iPSC microglia. **a)** Extended dose range transcriptomics for LY2090314 (9-point dilution series). Example concentration response curves for individual genes fitted to an exp5 model. For dose-response analyses, log-normalized gene expression was tested gene-wise with ANOVA for expression changes by dose; significant ANOVA genes (p < 0.05) with |log2FC| > 0.3 in any two doses were considered “dose-response genes” (n=1236 genes) and fitted to an exp5 model with non-linear least squares regression (see Methods, and Supplementary Table 15 for individual gene curve-fit parameters). **b)** Concentration-dependent pathway modulation by LY2090314. The dot-heatmap summarizes pathways significantly regulated across concentrations. The color of each dot represents median of the signed maximum |log2FC| across all dose-response genes within a given pathway. The dot size reflects the significance from overrepresentation analysis using a hypergeometric test after applying the Benjamini-Hochberg method for multiple hypothesis correction. The x-axis position of black-bordered and magenta-bordered dots reflect the median of the predicted benchmark dose (BMD, black) or EC50 (magenta) of the hits for given pathways (see Methods). **c)** Network view of selected pathways enriched by LY2090314. Graph layout of significantly enriched pathways highlights interconnected modules. Depicted nodes include dose-response selected genes belonging to each connected pathway. Color of each gene node represents its signed maximum |log2FC| from CTR on the doses tested.

Collectively, we have demonstrated that pharmacological modulation of the FAC-induced DLaM-like transcriptional state leads to corresponding functional changes in microglial phagocytosis, lipid handling, and lysosomal activity, with the reported GSK3 inhibitor LY2090314 as the most promising hit showing particularly broad restorative effects across transcriptional and functional readouts.

### LY20G0314 treatment induces a new microglia state characterized by lysosomal and lipid metabolic processes

We next performed scRNA-seq on FAC-stimulated hiPSC-derived microglia treated with or without LY2090314 to assess microglial state composition changes at single-cell resolution. UMAP of all microglial cells showed substantial overlap between samples while preserving clear cluster structure (Fig. 8a, small non-microglial clusters excluded, see Extended Data Fig. 10a-c). Unsupervised clustering identified multiple microglial populations, including DLaM15-like and Control, with LY2090314-treated samples giving rise to two new clusters which we called EnLyso/DLaM15-like, and EnLyso based on marker expression and pathway enrichment (Fig. 8b, Supplementary Table 17/18). The EnLyso cluster was enriched for lipid metabolic processes, lysosomal lumen, and vacuolar acidification, while the EnLyso/DLaM15-like cluster represented an intermediate state between DLaM15-like and EnLyso states. To evaluate treatment effects while correcting for technical variation, reciprocal PCA integration was applied to DMSO- and LY2090314-treated samples (Extended Data Fig. 10d). Integrated embeddings showed effective alignment across conditions, allowing direct comparison of cell state distributions. LY2090314 treatment altered the relative abundance of several clusters compared to untreated or FAC alone, including a reduction in the DLaM15-like population, an increase in the IFN-high population, and induction of the novel DLaM15-like/EnLyso and EnLyso-associated states (Fig. 8c).

**Fig. 8.**
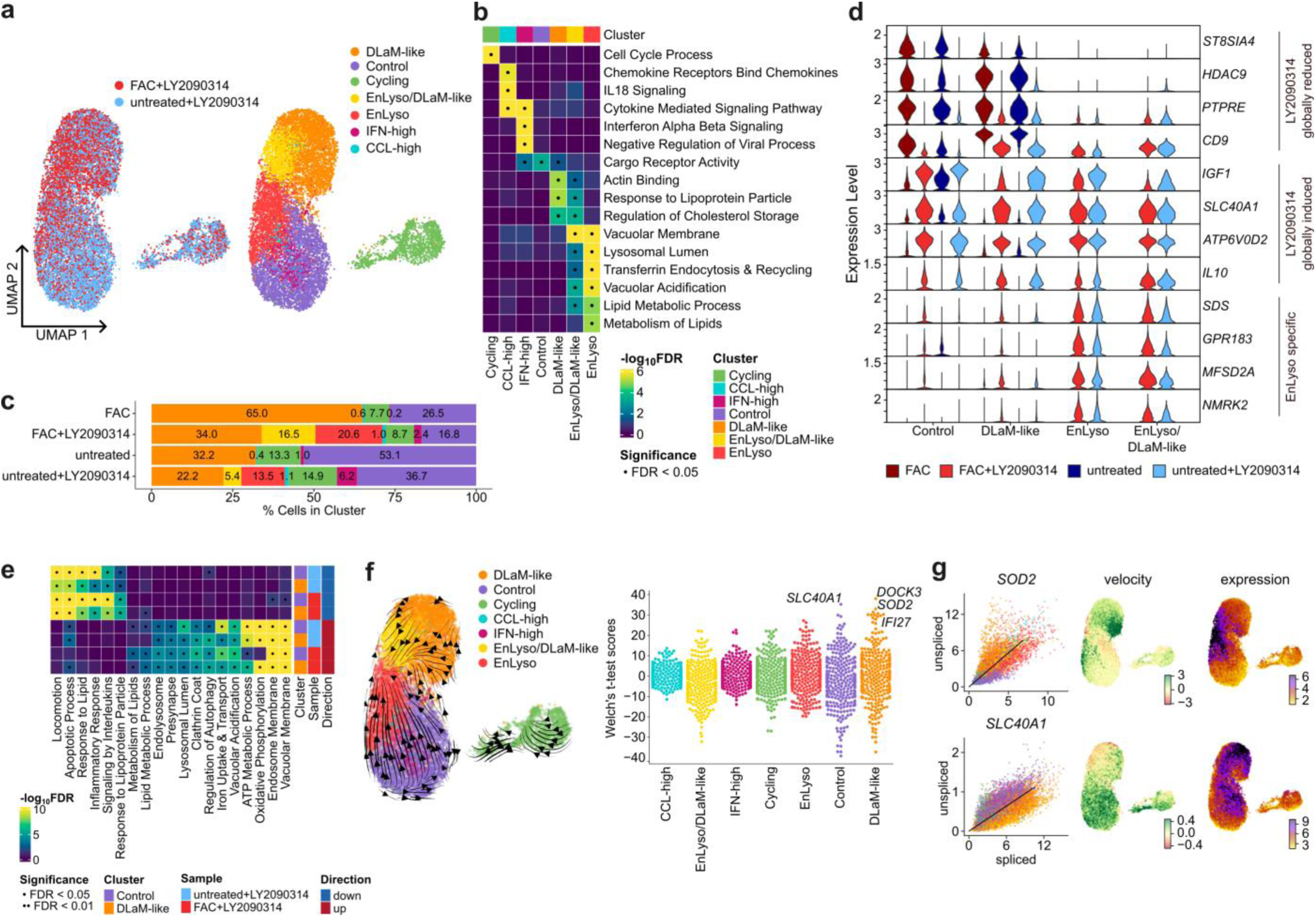
LY20G0314 treatment induces a microglial state enriched for lysosomal and lipid-metabolic programs. **a)** Single-cell sequencing results of LY2090314-treated hiPSC microglia. UMAPs of unstimulated and FAC-stimulated hiPSC microglia treated with 100 nM LY2090314 (left: colored by sample; right: by clusters). Unsupervised clustering recovers DLaM-like, Control-like, Cycling, IFN-high and CCL-high populations (already observed in Fig. 4) and reveals additional LY2090314-induced clusters EnLyso/DLaM-like and EnLyso. **b)** Heatmap of pathway enrichment by cluster. Enrichment was performed using a hypergeometric test including up-regulated DEGs in each cluster and applying the Benjamini-Hochberg method for multiple hypothesis correction. ((·) = FDR < 0.05). **c)** Stacked bars quantifying population proportions in the different hiPSC microglia samples. **d)** Treatment-responsive genes within selected clusters. Violin plots illustrate transcriptomic changes for selected genes induced by LY2090314 globally and specifically in the EnLyso cluster. **e)** Pathway-level response to LY2090314 across clusters. Enrichment was performed using a hypergeometric test and applying the Benjamini-Hochberg method for multiple hypothesis correction (· = FDR < 0.05). **f)** scRNA-seq velocity and trajectory analysis. Left: Visualizations on UMAP depict putative transitions between microglia states. Right: Scores from Welch t-test with overestimated variance are depicted with scores from individual genes in each population shown and selected genes in DLaM-like and Control populations highlighted. **g)** Scatter plots display quantifications of spliced and unspliced transcripts for selected genes with steady-state fits. Velocity estimates and smoothed expression for corresponding genes are shown on the UMAPs.

Differential expression analysis within clusters identified genes globally reduced or induced by LY2090314, as well as genes exhibiting cluster-specific regulation (Fig. 8d, Extended Data Fig. 10e, Supplementary Table 19). Markers associated with inflammatory activation, including *PTPRE*, *CDS*, and *HDACS* showed reduced expression across multiple clusters, whereas genes linked to lysosomal function and metabolic homeostasis, such as *ATP6V0D2*, *IL10,* and *IGF1*, were globally upregulated in all clusters following treatment with LY2090314. Several genes were selectively induced in EnLyso-associated states, like *GPR183* or *NMRK2.* Pathway-level analysis of genes altered by LY2090314 revealed coordinated transcriptional changes across clusters (Fig. 8e, Supplementary Table 20). Pathways related to lysosomal function, vacuolar membranes, endocytosis, lipid metabolism, and oxidative phosphorylation were significantly upregulated in all treated clusters and significantly enriched in EnLyso and EnLyso/DLaM15-like clusters. In contrast, inflammatory and cytokine signaling pathways as well as pathways related to locomotion and response to lipids showed broad downregulation across all clusters following treatment with LY2090314. Finally, we measured microglia cell state transitions using RNA velocity (Fig. 8f). This analysis revealed a convergent trajectory of cells into the EnLyso state originating from either the Control or DLaM15-like populations. In particular, the iron transporter, *SLC40A1*, exhibited a selective increase in RNA velocity within Control microglia while the antioxidant enzyme, *SOD2*, exhibited a marked increase in RNA velocity within the DLaM15-like population (Fig. 8g).

Together, these data demonstrate that LY2090314 reshapes microglial state composition and transcriptional programs at single-cell resolution, preferentially reducing disease- and lipid-associated signatures while promoting lysosomal and metabolic pathways.

## Discussion

Recent advances in single-cell transcriptomics have generated increasingly detailed cellular atlases of Alzheimer’s disease, revealing a complex landscape of microglial states associated with pathology and cognitive decline. However, how these microglia states contribute to disease, and whether modulating microglia identity in a patient represents a credible therapeutic approach, is still debatable. Here, we combined large-scale human single-nuclei transcriptomics with perturbational modeling and transcriptomic screening to establish a framework for identifying, reconstructing, and pharmacologically modulating Alzheimer’s disease-associated microglia states.

Our integrated analysis reinforces the emerging understanding that disease- and lipid-associated microglia represent a conserved endpoint of microglial remodeling in AD. Multiple independent studies have identified closely related microglia populations characterized either by expression of *APOE*, *TREM2*, *LPL* and other lipid-handling genes^3,6,20,24–27^ or functionally by altered lipid metabolism, metabolic stress adaptation and phagolysosomal remodeling^28–30^. Furthermore, our RNA velocity analysis revealed directional transitions from surveilling and intermediate states toward DLaM, supporting previous reports that disease-associated microglia emerge through progressive transcriptional remodeling rather than representing discrete lineages^27,31^. Together, these findings suggest that chronic AD-associated stress drives microglia toward metabolically specialized states marked by impaired lipid handling and lysosomal dysfunction.

Additionally, our data confirm that DLaM populations are heterogenous^6^. DLaM8 exhibited greater transcriptional plasticity, showed dynamic regulation of *ACSL1*, a key regulator of lipid-droplet-accumulating microglia^29^, overlapping with previously described plaque-associated microglia^6,32^. In contrast, DLaM15 exhibited a transcriptionally more constrained phenotype, was preferentially associated with tau pathology and cognitive decline^6^, and showed AD-associated transcriptional dynamics involving *CXCR4* and *LIPG*. Thus, we expanded on characterizing DLaM8 and DLaM15 as distinct stages within the disease-associated microglia continuum concordant with previously described DAM states^6^.

We identified FAC stimulation of hiPSC microglia as an i*n vitro* model to interrogate mechanisms capable of modulating DLaM15 microglia. Among a diverse panel of AD-relevant perturbations, FAC uniquely induced a robust DLaM15-like transcriptional state, characterized by expression of *APOE*, *TREM2*, *LPL*, *PPARG*, *MITF,* and *CD9*. Importantly, this signature showed greater concordance with human DLaM populations than previously reported *in vitro* models based on transcription factor engineering or epigenetic perturbation^7–9^. Single cell RNA-seq confirmed a marked expansion of a DLaM15-like population accompanied by depletion of surveilling microglia supporting the model where metabolic stress drives microglia from homeostatic toward disease-associated states. Functionally, FAC induced impaired Aβ handling, reduced lysosomal activity, and lipid accumulation indicating a broader failure of cellular clearance and metabolic adaptation and resembling disease-associated microglia observed in AD brains and human xenograft models, as well as foamy macrophages in atherosclerotic lesions^28,29,33,34^. These data establish FAC-treated hiPSC microglia as a disease-relevant model of human AD-associated microglia.

The ability of FAC to induce this phenotype is biologically plausible given the increasingly recognized role of iron-driven metabolic dysfunction in AD. Iron accumulates in amyloid plaques, dystrophic neurites and activates microglia during aging and disease, where chronic uptake of iron-rich cellular debris and Aβ aggregates promotes oxidative stress, lysosomal impairment and lipid metabolic dysfunction^35–39^. Consistently, FAC induced expression of ferroptosis-associated genes that are also markers of DLaM15, including *TXNRD1*, *PLA2G7* and *CCL3*, increased lipid droplet abundance and impaired lysosomal degradation. These observations align with reports demonstrating that reactive oxygen species are required for lipid droplet formation^40^ and that microglia are more susceptible to ferroptosis than other CNS cell types^41,42^. Together with the RNA velocity analysis linking Enhanced Redox microglia to DLaM15 through dynamic regulation of *ACSL1*, these findings suggest that chronic iron-induced metabolic stress is a major driver of disease-associated microglial remodeling, integrating oxidative stress, lipid dysregulation and lysosomal dysfunction into the transcriptional program characteristic of DLaM15.

A central advance of this study is the use of human disease-derived transcriptomic states as screening phenotypes. Most therapeutic discovery efforts in neurodegeneration focus on individual genes, pathways or functional readouts. In contrast, recent work in perturbational genomics and cell-state engineering has demonstrated the utility of high-dimensional transcriptional programs as quantitative phenotypes for drug discovery^13–16^. By using the DLaM15 signature as a screening endpoint, we were able to identify compounds that modulate an entire disease-associated cellular program rather than single genes or phenotypes. This strategy goes beyond disease signature-reversion approaches and provides a strategy for translating large-scale cellular atlases into experimental therapeutic discovery platforms. Indeed, deconvolution of comprehensive sets of patient states allowed not only to assess signature reversal, but was consistent with known mechanisms of action, and aided interpretation of compound-related effects.

A key finding of our screen is that transcriptional movement away from a disease-associated state does not necessarily indicate functional recovery. Instead, biological consequences depend on the state toward which DLaM15-like microglia are redirected. For instance, SYK and BTK inhibitors produced the strongest transcriptional alteration of the DLaM15-like signature, primarily shifting cells towards a DLaM8-like state, but both inhibitor classes also impaired Aβ phagocytosis. This could be explained by the concordance of DLaM8 and DLaM15 to the previously described Mic12 and Mic13 populations^6^, in which Mic12 is preferentially associated with amyloid pathology and reduced amyloid clearance, whereas Mic13 correlates with tau pathology and cognitive decline. Accordingly, the shift toward a DLaM8-like state may reflect redistribution within the disease-associated microglia continuum rather than restoration of a functionally competent phenotype. Similarly, TLR activators reduced the DLaM15 signature and improved Aβ phagocytosis, but did so through markedly inducing inflammatory signaling. Together, these findings emphasize that therapeutic modulation of microglia should be evaluated by integrating both transcriptomic and functional readouts with attention paid to the pharmacologically induced end-state.

Notably, we found that the previously described GSK3 inhibitor LY2090314 produced a fundamentally different outcome. Beyond reversing the DLaM15-like transcriptional program, LY2090314 induced a previously unrecognized lysosomal-metabolic state we termed EnLyso. This transcriptional transition was accompanied by restoration of lysosomal degradation, reduced lipid droplet accumulation, and improved Aβ phagocytosis. The molecular characteristics of EnLyso are consistent with previously described biological functions of GSK3 signaling, where GSK3 inhibition promotes TFEB activation, lysosomal biogenesis and autophagic flux^43–46^. Furthermore, GSK3 inhibition has been shown to improve pathology in AD mouse models through increased phagolysosome function and anti-inflammatory signaling^47,48^. Concordantly, LY2090314 induced *IL10* broadly, while reducing inflammatory markers such as *PTPRE* or *CDS*^49–52^. Finally, downregulation of *MITF* and *PPARG*, central regulators of DAM identity^7^ coordinating other lipid-associated genes, suggests that LY2090314 disrupts the transcriptional network program characteristic of DLaM15. Collectively, these findings suggest that LY2090314 relieves a central metabolic bottleneck linking iron overload, lipid dysregulation and impaired lysosomal function.

However, the beneficial effects of LY2090314 cannot be unequivocally attributed to inhibition of GSK3 alone. Although LY2090314 is among the most potent ATP-competitive GSK3 inhibitors reported^17^ showing effects in range within its described GSK3 potency, two additional chemically distinct GSK3 inhibitors produced markedly different transcriptional responses. The ATP-competitive inhibitor BIO-acetoxime^53^ induced partial features of the LY2090314 signature only at micromolar concentrations, whereas the non-ATP-competitive inhibitor Tideglusib^54^ showed little effect on microglial state. As allosteric kinase inhibitors exhibit broader kinase selectivity than ATP-competitive inhibitors^55^, these findings suggest that the LY2090314 effects may involve inhibition of an additional kinase target, or a combination of targets, besides GSK3 itself. Despite the uncertainty regarding the primary molecular target, the LY2090314 downstream cellular response was remarkably consistent, as treatment improved microglial function in all assessments. Altogether, these data support an emerging view that therapeutic benefit in AD is likely to arise from reprogramming disease-associated microglia toward functionally competent states rather than suppressing microglial activation.

Several limitations should be acknowledged. Although FAC recapitulates many features of DLaM15 microglia, disease-associated states *in vivo* are shaped by interactions with neurons, astrocytes, oligodendrocytes, and vascular cells absent in monoculture systems. Thus, while transcriptomic similarity between FAC-treated microglia and human DLaM15 was substantial, additional environmental cues are likely required to fully reproduce the disease-associated states observed in human brain. For instance, the effect of DLaM15 microglia on tau aggregation and subsequent neuronal dysfunction could be addressed in co-culture systems. Additionally, bioinformatic techniques to deconvolute cell type/state composition are limited by the decisions made on the signature selection and parameters of the calculation such as number of markers, so detected changes in cell state estimations must be experimentally verified. Future studies integrating multicellular systems, spatial transcriptomics and longitudinal perturbational analyses will be important for refining these models and validating the functional relevance of states such as EnLyso *in vivo*.

Overall, our study demonstrates that human disease-associated transcriptomic states can serve as experimentally tractable phenotypes for therapeutic discovery. By reconstructing a disease-associated microglial state directly from patient-derived transcriptomic data, using it as the basis for pharmacological screening, we identify pathways capable of reprogramming dysfunctional microglia toward a metabolically competent state. More broadly, these findings establish a general framework for translating increasingly comprehensive human cellular atlases into mechanism-based therapeutic discovery for neurodegenerative disease.

## Online Methods

### Human tissue collection

The tissues were obtained from The Netherlands Brain Bank (NBB, Netherlands Institute for Neuroscience, Amsterdam). All material has been collected from donors for or from whom a written informed consent for a brain autopsy and the use of the material and clinical information for research purposes had been obtained by the NBB. Samples were selected from the frontal and temporal cortex of individuals with Alzheimer’s disease (AD, n = 30) and age-matched cognitively normal controls (n = 42). Neuropathological assessment included evaluation of amyloid plaques and neurofibrillary tangles^56^. Clinical diagnoses were derived from clinical records. Individuals were assigned to joint diagnostic categories (CTR and AD; Table 1).

### Tissue processing and RNA quality control for NBB dataset

10 µm cryo sections were used to isolate RNA for quality control of patient samples. Tissue was lysed and homogenized using Trizol and chloroform in TissueLyser (QIAGEN) (2×10 min, 30 Hz). For phase separation, samples were centrifuged (3000 rpm, 10 min, RT). RNA was purified by isopropanol precipitation and AMPure XP beads purification. RNA quantity was assessed by Quant-iT^TM^ RNA assay kit (Thermo Fisher Scientific). RNA integrity (RIN) was measured by Bioanalyzer 2100 Expert (Agilent) using RNA 6000 Pico Kit.

### Single-nuclei RNA library preparation, sequencing and primary preprocessing of NBB samples

Nuclei were isolated in accordance with the 10x Genomics guidelines for single-cell RNA-seq sample preparation (protocol CG000124). Briefly, 25 μm cryosections from frozen tissue were collected into 1.4 mL tubes (Micronic) and homogenized in 800 μL nuclei extraction buffer (Miltenyi) using 5 mm metal beads (2 × 30 s at 25 Hz; TissueLyser, (QIAGEN)). Homogenates were passed through a 100 μm cell strainer (Merck), and nuclei were pelleted by centrifugation (500 × g, 5 min, 4 °C). Pellets were washed in 800 μL wash buffer (10 mM Tris-HCl, pH 7.4; 10 mM NaCl; 3 mM MgCl₂; 0.5 U/μL RNase inhibitor; 1% BSA; 1 mM DTT). To remove debris and sample-specific contaminants, nuclei were further purified by density gradient centrifugation. Nuclei suspended in 200 μL wash buffer were underlaid with 600 μL gradient buffer (30% (w/v) OptiPrep; 10 mM Tris-HCl, pH 7.4; 10 mM NaCl; 3 mM MgCl₂; 0.2 U/μL RNase inhibitor; 1% BSA; 1 mM DTT) and centrifuged (4,000 × g, 15 min, 4 °C). The nuclei pellet was subjected to two additional wash steps (800 μL wash buffer; 500 × g, 5 min, 4 °C) and filtered through a 40 μm strainer (Merck). Nuclei were quantified using an automated cell counter (Luna FX7, Logos Biosystems) with AO/PI staining, and concentrations were adjusted to target 20,000 nuclei per sample. Single-cell 3′ gene expression libraries were generated using the Chromium Next GEM Single Cell 3′ Reagent Kit v3.1 (10x Genomics) according to the manufacturer’s instructions (protocol), with Agencourt AMPure XP beads used for all clean-up steps. Full-length cDNA was amplified by PCR, purified and quantified using the Qubit dsDNA High Sensitivity assay (Thermo Fisher Scientific). Fragment size distributions were assessed using the Agilent TapeStation High Sensitivity D5000 assay, evaluating fragments between approximately 200 and 9,000 bp. Sequencing libraries were then constructed from amplified cDNA following the Chromium Next GEM 3′ v3.1 protocol. Final libraries were assessed on the Agilent TapeStation using the DNA 1000 or HS D5000 assay to determine average fragment size (typically ∼470–550 bp). Libraries were pooled equimolarly and sequenced using an Illumina NovaSeq 6000 S4 platform to a depth of at least 20,000 reads per nucleus. Sequencing reads were aligned to the human reference genome (GRCh38) with GENCODE v32 annotation using Cell Ranger (v7.0.0) with intronic reads included (--include-introns). Barcodes identified as cell-containing droplets by Cell Ranger were retained for downstream analysis.

### Generation of human metacell reference atlas

A previously published human brain single-cell atlas^19^ comprising 606 samples in *loom* format was used to construct a metacell reference atlas. Metacells were generated independently for each sample using metacells-2 (v0.8.0). Purity was assessed using the authors’ 31-cluster annotation, with 121,097 metacells (97%) classified as pure. Raw counts from pure metacells were log-normalized and scaled, highly variable genes were selected, and principal component analysis was performed (50 PCs). Batch correction was applied using *Harmony* with donor as a covariate, followed by UMAP embedding (return.model = TRUE).

### Public data integration and metadata harmonization

Public datasets from ROSMAP^6^, SEA-AD^57^ and Morabito et al.,^58^ were obtained from Synapse repositories (syn31512863, syn26223298, syn26670419), while remaining public datasets^31,59,60^ were retrieved from Gene Expression Omnibus (GSE167494, GSE157827, GSE148822). FASTQ files were downloaded for all datasets except Gerrits et al., for which BAM files were converted to FASTQ files using the *bamtofastq* tool within Cell Ranger (v7.0.0)^61^. All datasets were reprocessed using the same preprocessing pipeline as the newly generated data. For pooled ROSMAP libraries, cells were assigned to donors based on published demultiplexed barcodes. Clinical and neuropathological metadata, including diagnosis, APOE ε4 status, age, sex, and brain region, were curated from original publications and associated metadata files. Metadata fields were harmonized across studies to define comparable diagnostic categories (Table 1).

### Downstream snRNA-Seq analyses

Filtered count matrices from Cell Ranger were imported into R (Seurat) on a per-sample basis, normalized, and genes subset to variable features in the metacell reference atlas. Samples were mapped to the reference atlas using Azimuth^62,63^ via Seurat (*FindTransferAnchors* and *MapQuery*, enabling projection onto the reference UMAP and transfer of cell type annotations with associated confidence scores. Nuclei assigned to the same cell type were aggregated across samples to generate cell type-specific objects. Cell type-specific datasets were integrated using Scanpy^64^ including log-normalization, scaling, principal component analysis, and Harmony batch correction with study and sex as covariates. Clustering was performed using the Leiden algorithm at multiple resolutions, followed by UMAP visualization. Doublets and low-quality nuclei were removed through iterative clustering and filtering. Final annotations were assigned based on previously published canonical marker genes from snRNA-Seq datasets^4,6,57^, yielding a dataset comprising 3.6 million nuclei with harmonized cell type annotations across all public datasets and the newly generated NBB snRNA-Seq dataset.

### MAGMA analysis

Alzheimer’s disease GWAS summary statistics for the European population^1^ were integrated with single-cell expression data to identify cell type–specific genetic enrichment. SNP-to-gene mapping was performed using the MAGMA (Multi-marker Analysis of GenoMic Annotation, v1.10) pipeline^65^, accounting for linkage disequilibrium based on the 1000 Genomes European reference panel, with a 35 kb upstream and 10 kb downstream window. Gene-level specificity values were subsequently used as input for cell type enrichment analysis implemented in the R package MAGMA. Celltyping (v2.0.15). For each annotated cell type, genes were ranked according to their specificity metrics, and the 10% most specific genes were selected to define cell type-specific gene sets. Gene-level association statistics were then tested for enrichment within each cell type-specific gene set using the MAGMA gene-set analysis framework.

### Microglia subclustering

Nuclei annotated as microglia were subset and integrated into a dedicated object using the standard Scanpy workflow described above. Clustering using the Leiden algorithm (resolution = 1.2) identified 19 microglial clusters.

### Differential gene expression and marker identification

Cluster-specific marker genes were identified using a pseudobulk approach. Cells belonging to the same cluster within each sample were aggregated, and gene counts were summed. Pseudobulk samples with fewer than 10 cells were excluded. Cluster markers were calculated using the limma-voom framework, incorporating study and sex as covariates (*∼Cluster + Study + Sex*). Counts were transformed to log2 counts per million (CPM) and mean-variance relationships per gene were estimated using the *voom* function, followed by *lmfit* (weighted least square regression), where precision weights were determined by the *voom* fit. This was followed by *contrasts.fit* to obtain coefficients (log_2_FC estimates) and standard errors from the linear model fit for each of the pairwise comparisons (each microglia cluster against all other clusters), and finally *eBayes* to compute moderated t-statistics and F-statistics by empirical Bayes moderation of the standard errors. Disease-associated differential expression was similarly assessed at the pseudobulk level within each cell type (∼Diagnosis + Study + Sex). Disease-dependent dysregulated genes were then calculated using the limma-voom pipeline.

### Pathway/ontology enrichment and trait-association

Cluster markers genes (log_2_FC > 0.8, FDR < 0.05) were subjected to overrepresentation analysis (ORA) using the *enricher* function from the *clusterProfiler* library in R. GO terms, Wikipathways, Reactome and Hallmark DB were downloaded from MSigDB ^66^ and used in clusterProfiler. P-values were then corrected for multiple hypotheses testing using the Benjamini-Hochberg procedure for each set of pathways in each database separately. Pathway enrichment analysis was also performed on AD-associated DEGs (AD vs. CTR, FDR < 0.05) for each cell type. AD risk genes from Bellenguez et al.^1^ and enrichment was assessed using a hypergeometric test with Benjamini– Hochberg FDR correction.

### Differential abundance analysis

Differential abundance analysis was performed on single-nuclei microglia using a negative binomial generalized linear model with the *edgeR* package ^67^. Differences in abundance were estimated using empirical Bayes quasi-likelihood F-tests with the function *glmQLFTest* and Benjamini-Hochberg-corrected p-values. For the differential abundance analysis, cells from the proliferating and Surv_18 cluster (low abundance) were removed. Differential abundance was calculated for Joint Diagnosis, Clinical Diagnosis, CERAD score, Braak Stage, and AT score by comparing each disease category against its corresponding control group. For the AT score, A+ was defined as CERAD score of C1-C3, T+ samples had a Braak Stage ≥ 3, while A- and T-corresponded to CERAD C0 and Braak Stage <3, respectively.

### Velocity and Trajectory analysis

For RNA velocity analysis, coordinate-sorted BAM files from the NBB dataset were processed with velocyto (v0.17.17) using GRCh38 and repeat annotation from UCSC genome browser. Sample-specific loom files were integrated into a single h5ad and analyzed with scVelo (v0.3.3) ^68^ to recover dynamics, infer velocities, construct velocity graphs and estimate latent time. Differential velocity-associated genes were identified using scVelo’s *rank_velocity_genes* function, separately for AD and CTR samples.

### Microglia cluster similarities across studies

Cluster similarities were measured using the Jaccard index, defined as

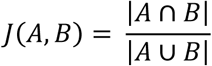

where A and B are defined as different sets of marker genes. Cluster markers for different microglia populations were derived from the supplemental tables of published studies ^3,6^, and the positive marker genes were compared to the microglia cluster markers from this work (log_2_FC > 0.58, FDR < 0.05).

### Cultivation and microglia differentiation of human iPSCs

To ensure uniformity and high quality of starting cells, cryopreserved single cell hiPSCs batches were created using the CryoPause method^69^, with minor modifications. Frozen hiPSCs were thawed at 37°C in a water bath for two minutes before transferred to a 50 mL tube containing 10 mL Essential 8™ (E8) Medium (ThermoFisher Scientific, #A1517001), then centrifuged for five minutes at 400 x g to create cell pellets. The supernatant was aspirated and the pellet resuspended in 10 mL E8 medium supplemented with 10 µM Y-27632 dihydrochloride (ROCK inhibitor) (BioTechne, #1254/10). We counted cells using a Chemometic NucleoCounter via resuspending 1 µL Solution 18 (Chemometic, #910-3018) in 20 µL of cell suspension, with 12 µL of the dyed cell solution added into the chambers of an NC-Slide A8 (Chemometic, #90279000). Cells were plated at a density of 5k - 30k cells/cm^2^ in E8 containing 10 µM Y-27632 dihydrochloride and incubated at 37°C and 5% CO_2_. Medium was exchanged after 24 hours to E8 without ROCK inhibitor, followed by a daily medium exchange. After culturing the cells for five days (to a confluency ∼50 −70%), cells were split using 0.5 mM EDTA (Sigma-Aldrich #03690) / 0.9% NaCl (Sigma-Aldrich #S5150) and re-seeded at a density of 5k - 12k cells/cm^2^. Cells were split every three to four days, as described above. hiPSCs were differentiated into microglia as previously described^70^, with minor modifications.

### In vitro perturbation experiments for AD model identification

*In vitro* perturbation experiments were conducted using a robotic liquid-handling workflow (Viaflo, Integra). Cryopreserved hiPSC-microglia were thawed and seeded into 384-well plates in standard cultivation medium and maintained at 37°C and 5% CO_2_. After attachment, cells were exposed to following treatments: aggregated amyloid beta 1-42 (AnaSpec #AS-20276, also see Section “Aβ phagocytosis in hiPSC microglia”), FAC (Sigma #F5879), neuraminidase (Sigma #1158588601), rotenone (SelleckChem #S2348), lipopolysaccharide from escherichia coli O55:B5 (Sigma #L5418) combined with interferon gamma (Immunotools #11343534), 2-deoxy-d-glucose (Sigma #D6134), catalytic domain of MMP-3 (Sigma #444217), CGS-21680 hydrochloride hydrate (Sigma #C141), Bleomycin sulfate (Abcam #ab142977), C75 (Tocris #2489), SC-26196 (Sigma #PZ0176) and corresponding buffer controls. Treatments were applied for up to 96h under standard culture conditions. Following stimulation, cells were washed and lysed directly on the assay plate. Lysates were sealed, snap-frozen on dry ice, and stored at −80°C for transcriptomic analysis.

### High-throughput transcriptomic screen in hiPSC microglia

#### Compound treatment

Cryopreserved hiPSC-microglia were thawed, recovered and seeded onto 384-well plates in standard cultivation medium, followed by incubation at 37°C and 5% CO_2_. Cells were subsequently exposed to 0.5 mM FAC (Sigma #F5879) or control medium for 48h before the addition of compounds (see Supplementary Table 12) via robotic liquid handling system for additional 24h. Cells were washed and lysed directly on the assay plate. Lysates were sealed, snap-frozen on dry ice and stored at −80°C.

#### High-throughput transcriptomic profiling (ScreenSeq)

High-throughput transcriptomics was processed using Evotec’s plate-based ScreenSeq^TM^ automated platform as previously described^22^. Following cell lysis, mRNA was labelled with a unique molecular identifier (UMI) on a well-basis. After pooled cDNA synthesis, adapters containing Illumina’s unique *dual indexes* were attached for final library preparation and sequenced on a NovaSeq 6000 system. Transcriptomic profiling was performed in batches of 384 maximum samples per plate. Stimulations (n = 3) and controls (n = 9) originated from the same plate (Fig. 3a). Reads were mapped to the human genome (GRCh38.p13 – accession GCA_000001405.28) using Spliced Transcripts Alignment to a Reference (STAR) (version 2.7.9a). Multi-mapped reads, reads with mapping to intergenic regions (with respect to Ensembl gene annotation information version 104) and reads with ambiguous mapping location (i.e. location associated with more than one gene) were discarded, and UMI-based deduplication was performed. Log-scaling of the resulting counts used CP10K (counts per 10 thousand) as scaling and 1 as pseudo-count unless indicated otherwise.

#### Dimension reduction visualizations

Transcriptomic data were visualized using a PCA on the 2000 most variable genes from the log-normalized count matrix with the *prcomp* function in R. For the FAC-reversal compound screen, UMAPs (2000 most variable features and a PCA space of 50 dimensions) were generated using the uwot package on ComBat-corrected (by screening plate; R package sva) log-normalized expression data.

#### Differential expression analysis and pathway enrichment

Pair-wise comparisons for microglia perturbations were performed against untreated microglia controls using the limma-voom pipeline. Compound effects were compared against FAC-only treated controls using limma-voom, with formula *∼Compound Dose + Plate* using Compound Dose as the contrast factor. Differentially expressed genes (DEGs) were calculated separately for every compound at every concentration. DEGs were subjected to pathway enrichment analysis as described above.

#### Cell type deconvolution analyses

To determine which stimulation in hiPSC microglia was best associated with patient-derived microglia states, the R package BRETIGEA (BRain cEll Type specIfic Gene Expression Analysis, version 1.0.3) ^23^ was applied. Normalized Screen-seq (bulk) count matrices derived from stimulated and/or compound-treated hiPSC-microglia were used as input for the *findCells* function with default parameters (nMarkers = 50, method=”SVD”). Deconvolution was performed against patient microglia cluster markers (FDR < 0.05 and log_2_FC > 0.3). Deconvolution scores are normalized relative to untreated hiPSC-derived microglia (Z-scored) and represent standard deviations from the control mean.

#### Microglia cluster and stimulation signature similarities

Similarities between microglia clusters and hiPSC microglia stimulation protocols were measured using a modified Jaccard index, defined as

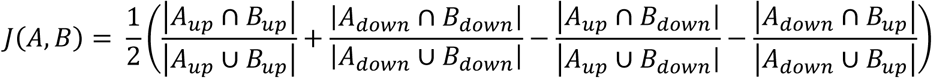

where *A_up_* and *A_down_* are defined as the set of genes up- or down-regulated in a microglia cluster compared to other clusters, and *B_up_* and *B_down_* are defined as the set of genes up-or down-regulated by a stimulus in hiPSC microglia. The modified Jaccard has values between −1 and 1, with positive values indicating a positive or shared overlap in the signature and negative values indicating a negative or reversed overlap in the signature.

A signature reversal (sigmoid) score was used to quantify the degree to which a compound was able to revert the FAC-induced transcriptomic signature. The sigmoid score was defined as:

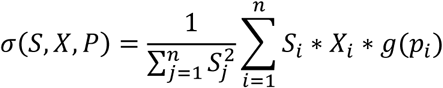

where *S* represent the FAC signature (log_2_FC estimate for all *n* DEGs (|log_2_FC| > 0.58; FDR <0.05) comparing FAC against CTR), *X* represents the log_2_FC estimates and *P* represents the adjusted p-value estimates for *n* DEGs induced by compound treatment (against FAC), and the function *g*(*y*) a sigmoid-like function mapping adjusted p-value estimates *p_i_* from values 0 to 1 to values 1 to 0 (*g*(0) = 1, *g*(1) = 0). Thus, a compound reverting the FAC-induced signature has a negative signature score, and a compound exacerbating it has a positive score.

#### Dose response analysis

The R package DRomics ^71^ was applied for dose-response fitting of compound-treated hiPSC microglia. For DEG number fit of compound reversal screen we obtained replicate-free DEG numbers per sample/well grouped by compound and concentration (n_Cpd=3, n_CTR=24), and inputted to the function *drcfit* with the custom exponential model 5 (“exp5” from the “Benchmark Dose Technical Guidance”, Risk Assessment Forum U.S. Environmental Protection Agency, https://www.epa.gov/risk/benchmark-dose-technical-guidance (2012)):

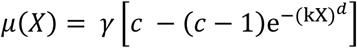

where *μ(X)* is the mean response at dose *X* > 0, γ is the background of normalized expression levels (0 DEGs for concentration 0), *c* is the value of the asymptote for the absolute maximum fitted deviation of the baseline, *k* is the slope, and *d* is the power. Because of the hill shaped curves obtained on the fitting, the EC50 can be obtained by applying the inverse function at *μ*(*X*)*_EC_*_50_ defined by:

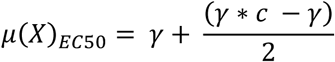

For the extended dose-range experiment (9 doses) of LY2090314, we followed the standard DRomics pipeline for individual gene fits. Briefly, normalized expression data was used for feature selection based on ANOVA and expression changes (FDR < 0.05, |log_2_FC| > 0.3 in at least two concentration points vs baseline); followed by model fitting to “exp5” as described above for each dose-responsive gene; finally *bmdcalc* was used to obtain benchmark doses based on 1 SD from predicted baseline, and dose responsive genes with BMD outside of the dose range (0-10μM) were filtered out (for curve parameters see Supplementary Table 15). The set of dose-response genes was used for enrichment analyses using the *enricher* function from the *clusterProfiler* library in R, as described above.

### Single-cell RNA-seq of hiPSC microglia

#### Experimental setup

Cryopreserved hiPSC-derived microglia were thawed and seeded into 24-well plates in standard cultivation medium and maintained at 37°C and 5% CO₂. After attachment, cells were treated with 0.5 mM FAC (Sigma, #F5879) or cultured in medium alone for 48 h, followed by addition of 100 nM LY2090314 (MedChemExpress #HY-16294) and incubation for an additional 24 h under standard conditions. For preparation of single-cell suspensions, cells were washed with PBS-/- (ThermoFisher Scientific #10010023), incubated with TrypLE Express (Gibco, #12604021) for 10 min at 37°C and 5% CO₂, and detachment was stopped with 0.04% BSA (Sigma #A8806). After one additional washing step, cells were ready as single-cell suspensions.

#### Single-cell suspension, RNA library preparation, and sequencing

Cells were prepared as single-cell suspensions with a viability greater than 90%. Cell suspensions were kept on ice and processed within 30 minutes of preparation. Cells were diluted to a final concentration of 1,000 cells/µL in nuclease-free water, according to the targeted recovery and expected capture efficiency. Cell numbers were determined prior to loading, and samples were adjusted individually to the required volume. RNA libraries were prepared and sequenced as described above.

#### Preprocessing and quality control

Raw reads were mapped to human reference genome version GRCh38 and the Gencode reference annotation (GENCODE v32) using the Cell Ranger (v7.0.0) pipeline with the --include--introns parameter. Filtered barcodes annotated as cell-containing droplets by Cell Ranger were extracted. Barcodes with percent mitochondrial UMIs greater than 15% or less than 3% were filtered for downstream analyses, as well as barcodes with number of UMIs less than 10^3.5^.

#### Normalization, clustering and cell type annotation

Raw counts from all samples were split initially into non-compound (DMSO+FAC / DMSO+untreated) and LY2090314 (Cpd) treated (Cpd+FAC / Cpd+untreated) to be analyzed independently before integration. Both sets were normalized, variable features detected, and normalized expressed values scaled. PCA was performed with 50 PCs followed by *harmony* batch-correction using sample as a covariate, then UMAP dimension reduction and Louvain clustering were applied.

For the non-compound set 5 clusters were obtained. Based on marker gene expression, 3 major clusters were annotated as microglia and 2 smaller clusters annotated as mesenchyme and mast cells were discarded (Extended Data Figure 5). Following this, microglial exclusive clustering was reapplied and identified clusters were annotated based on relative expression of marker genes (Control – P2RY6, IGF1, F13A1; DLaM15-like – LPL, CD9, TPRG1, PPARG, ITGAX, TREM2, MITF; Cycling – MKI67, TOP2A, CENPF; IFN-high – IFIT1, RSAD2, MX2; CCL-high – CCL4, CXCL8, RELB). Module scores were measured for marker genes of patient-data derived microglia clusters (log_2_FC > 0.58, FDR < 0.05) using Seurat’s *AddModuleScore* function.

On the other hand, clustering of the LY2090314 treated microglia samples yielded clusters with similar relative expression profiles as the untreated, with 2 additional clusters which were annotated as EnLyso, expressing high levels of SDS, MFSD2A, SDSL, NMRK2, and EnLyso/DLaM15-Like, which co-expressed markers of the EnLyso and DLaM15-like clusters.

Reciprocal PCA using Seurat’s *IntegrateData* function was applied to project each dataset into the PCA space of another dataset, after splitting the object by sample (four in total: DMSO+untreated, DMSO+FAC, Cpd+untreated, and Cpd+FAC).

#### Differential expression and pathway enrichment

Cluster markers for different hiPSC microglia cell types in samples treated with LY2090314 were estimated using a Wilcoxon rank sum test, with p-values corrected for multiple hypothesis testing using the Benjamini–Hochberg procedure. Cluster markers for each microglia cluster were extracted (log_2_FC > 0.15, FDR < 0.05) and pathway enrichment analysis was performed using the *enricher* function from the clusterProfiler library in R with pathways downloaded from MSigDB and correction for multiple hypothesis testing using the Benjamini-Hochberg procedure.

Differentially expressed genes were obtained by comparing the LY2090314 treated samples (Cpd) against non-treated (DMSO), within the FAC and untreated samples paradigm, and in the two major populations detected – Control and DLaM15-like, using a Wilcoxon rank sum test, with p-values corrected for multiple hypothesis testing using the Benjamini–Hochberg procedure. This resulted in four comparisons: Cpd+FAC vs. DMSO+FAC in Control microglia, Cpd+FAC vs. DMSO+FAC in DLaM15-like microglia, Cpd+untreated vs. DMSO+untreated in Control microglia, and Cpd+untreated vs. DMSO+untreated in DLaM15-like microglia. Genes that were up-regulated in all four comparisons (log_2_FC > 0.15, FDR < 0.05) were annotated as globally induced by LY2090314, and those that were down-regulated in all four comparisons (log_2_FC < −0.15, FDR < 0.05) were annotated as globally reduced by LY2090314. EnLyso marker genes that were not globally induced by LY2090314 were annotated as EnLyso specific.

Pathway enrichment analysis was performed across the four comparisons described above using the *enricher* function from the clusterProfiler library in R with pathways downloaded from MSigDB and correction for multiple hypothesis testing using the Benjamini-Hochberg procedure.

#### Velocity analysis

RNA velocity was performed on LY2090314 treated samples with velocyto (v0.17.17) using coordinate-sorted bam files as input. Then, sample-specific loom output files generated by velocyto were integrated into one h5ad file that was used as an input for scVelo (v0.3.3). Using scVelo, velocity analysis was performed using the deterministic mode and velocity embeddings were estimated. A differential expression test was performed using scVelo’s *rank_velocity_genes* function which employs a Welch’s t-test with overestimated variances on the velocity expression values to find genes that are differentially transcriptionally regulated in one cluster compared to all other clusters.

#### Lipid droplet quantification in hiPSC microglia

Neutral lipids were labeled with BODIPY™ 493/503 (ThermoFisher Scientifc #D3922) according to manufacturer’s instructions with minor changes. In short, cryopreserved microglia were plated onto 384-well plates in standard cultivation medium and, following attachment, either treated with 0.5 mM FAC (Sigma # F5879) or maintained in culture medium alone for 48h at 37°C and 5% CO_2_. Where appropriate for evaluation of LY2090314’s effects, the compound was subsequently added in the concentrations indicated in Fig. 6, and cells were incubated for an additional 24h at 37°C and 5% CO_2_ prior to fixation with 4% paraformaldehyde (PFA; Science Services GmbH #E15713-S) and BODIPY™493/503 plus nuclear counter (Hoechst 33258; ThermoFisher #H3569) staining. Fixed cells were imaged using an Opera Phenix high-content confocal microscope (Revvity), and lipid droplet area was quantified from BODIPY™ 493/503-positive cytoplasmatic structures in the Alexa Flour 488 channel using Harmony image analysis software (Revvity).

#### Lysosomal activity (DQ-BSA) assay in hiPSC microglia

Lysosomal proteolytic activity in hiPSC-microglia was determined using DQ™-red BSA (ThermoFisher Scientifc #D12051). Matching DQ™-red BSA concentration between conditions (also see Fig. 4f and Extended Data Fig. 5d) were identified using Alexa Fluor™ 488 conjugate-BSA (ThermoFisher Scientific #A13100) titration. Cryopreserved microglia were thawed and plated onto 384-well plates in standard cultivation medium. Following attachment, cells were either treated with 0.5 mM FAC (Sigma #F5879) or cultured in medium alone for 48h at 37°C and 5% CO2, after which LY2090314 was added (where appropriate, see Fig. 6) and incubation continued for 24h at 37°C and 5% CO_2_. Prior to live-cell imaging, cells were pre-incubated for 1h with 3x concentrated DQ-BSA or Alexa Fluor™ 488 conjugate-BSA, followed by 1h incubation with nuclear counterstaining (Hoechst 33258; ThermoFisher #H3569). After a washout step, original treatment conditions were reapplied, and live-cell imaging was initiated immediately performed on an Operetta high-content imaging system (Revvity), with images acquired hourly for up to 24h. Matching BSA loading concentration per condition were identified by quantification of intracellular Alexa Fluor™ 488 conjugate-BSA fluorescence spot intensity in the Alexa Flour 488 channel using Harmony image analysis software. Proteolytic activity was quantified from intracellular DQ™-red BSA fluorescence spot intensity in the Alexa Fluor 594 channel using Harmony image analysis software.

#### Aβ phagocytosis in hiPSC microglia

Amyloid-β(Aβ)_1-42_ was prepared as aggregated material for phagocytosis assays. Briefly, Aβpeptides (Anaspec #AS-20276) were dissolved in DMSO to generate monomeric stock solutions, aliquoted, and stored at −80°C until usage. For aggregation, monomers were diluted in NSP buffer (10nM potassium phosphate, 10mM NaCl, pH 7.4) and incubated at 37°C under controlled agitation to generate Aβ-aggregated species. Following aggregation, samples were subjected to controlled sonication to obtain uniformly dispersed aggregated and stored at −80°C.

Aggregated Aβ was labelled with pHrodo™ Green STP ester (ThermoFisher Scientific #P35369) according to a protocol adapted from Fujifilm/Cellular Dynamics^72^. In brief, aggregates were buffer exchanged into bicarbonate labeling conditions, incubated with pHrodo dye for 1.5h, and subsequently washed extensively to remove unbound dye. Labelled Aβ aggregates were resuspended in Hank’s Balanced Salt Solution (ThermoFisher Scientific #14185052) and stored at −80°C until usage.

Aβ uptake by hiPSC-derived microglia was assessed using the pHrodo-labelled Aβ-aggregates. Cryopreserved hiPSC-derived microglia were thawed and seeded onto 384-well plates in standard cultivation medium, followed by incubation at 37°C and 5% CO_2_. For stimulation, cells were exposed to 0.5mM FAC for 24h, followed by the addition of compounds (where appropriate, also see Fig.6 and Supplementary Table 12), and incubation continued for further 24h at 37°C and 5% CO2. Subsequently, pHrodo-labelled Aβ-aggregates together with nuclei counterstain NucLight Rapid Red (Sartorius #4717) were added, and live-cell imaging was performed on an Incucyte SX5 system (Sartorius) or for single cell analysis on an Operetta high-content imaging system. Images were acquired hourly for up to 24h at 37°C and 5% CO_2_. Aβuptake was quantified based on intracellular pH-dependent fluorescence intensity. For Incucyte-based analysis, total integrated intensity in the green channel was normalized to the nuclei count per well detected in the orange channel. For Operetta-based analysis, Aβ uptake was quantified as spot intensity of the Alexa Flour 488 channel and normalized to the nuclear signal detected in the Alexa Flour 597.

## References

1. Bellenguez, C. et al. New insights into the genetic etiology of Alzheimer’s disease and related dementias. Nat. Genet. 54, 412–436 (2022).

2. Lambert, J. C. et al. Consensus meta-analysis of genome-wide association studies for Alzheimer’s disease and related dementias. Nature Genetics Preprint at 10.1038/s41588-026-02583-1 (2026).

3. Sun, N. et al. Human microglial state dynamics in Alzheimer’s disease progression. Cell 186, 4386–4403.e29 (2023).

4. Mathys, H. et al. Single-cell transcriptomic analysis of Alzheimer’s disease. Nature 570, 332–337 (2019).

5. Prater, K. E. et al. Human microglia show unique transcriptional changes in Alzheimer’s disease. *Nat*. Aging 3, 894–907 (2023).

6. Green, G. S. et al. Cellular communities reveal trajectories of brain ageing and Alzheimer’s disease. Nature 633, 634–645 (2024).

7. Dolan, M. J. et al. Exposure of iPSC-derived human microglia to brain substrates enables the generation and manipulation of diverse transcriptional states in vitro. Nat. Immunol. 24, 1382–1390 (2023).

8. Haage, V. et al. HDAC inhibitors engage MITF and the disease-associated microglia signature to enhance amyloid β uptake. Brain Behav. Immun. 129, 279–293 (2025).

9. Podleśny-Drabiniok, A. et al. BHLHE40/41 regulate microglia and peripheral macrophage responses associated with Alzheimer’s disease and other disorders of lipid-rich tissues. Nat. Commun. 15, (2024).

10. Haghighi, M., Caicedo, J. C., Cimini, B. A., Carpenter, A. E. & Singh, S. High-dimensional gene expression and morphology profiles of cells across 28,000 genetic and chemical perturbations. Nat. Methods 19, 1550–1557 (2022).

11. Kopylova, I. V. et al. Convergent transcriptomic signature in iPSC-dopaminergic neurons of hereditary Parkinson’s disease. Life Sci. Alliance 9, (2026).

12. He, C. et al. Integrating population-level and cell-based signatures for drug repositioning. (2025) doi:10.5281/zenodo.16791909.

13. Wilson, D. M., et al. Hallmarks of neurodegenerative diseases. Cell vol. 186 693–714 Preprint at 10.1016/j.cell.2022.12.032 (2023).

14. Guin, D. et al. High-throughput transcriptomic screening reveals entrectinib as a repositioning opportunity in 19q12 autism spectrum disorder. Sci. Rep. 15, (2025).

15. Fisher, J. L. et al. Signature reversion of three disease-associated gene signatures prioritizes cancer drug repurposing candidates. FEBS Open Bio 14, 803–830 (2024).

16. Caldwell, A. B. et al. Endotype reversal as a novel strategy for screening drugs targeting familial Alzheimer’s disease. Alzheimer’s and Dementia 18, 2117–2130 (2022).

17. Atkinson, J. M. et al. Activating the Wnt/β-catenin pathway for the treatment of melanoma - Application of LY2090314, a Novel selective inhibitor of glycogen synthase kinase-3. PLoS One 10, (2015).

18. Martinkova, J., et al. Proportion of Women and Reporting of Outcomes by Sex in Clinical Trials for Alzheimer Disease A Systematic Review and Meta-analysis. JAMA Netw. Open 4, E2124124 (2021).

19. Siletti, K. et al. Transcriptomic diversity of cell types across the adult human brain. Science (1979). 382, (2023).

20. Srinivasan, K. et al. Alzheimer’s Patient Microglia Exhibit Enhanced Aging and Unique Transcriptional Activation. Cell Rep. 31, (2020).

21. Srinivasan, K. et al. Alzheimer’s Patient Microglia Exhibit Enhanced Aging and Unique Transcriptional Activation. Cell Rep. 31, (2020).

22. Rosell-Hidalgo, A. et al. In-depth mechanistic analysis including high-throughput RNA sequencing in the prediction of functional and structural cardiotoxicants using hiPSC cardiomyocytes. Expert Opin. Drug Metab. Toxicol. 20, 685–707 (2024).

23. McKenzie, A. T. et al. Brain Cell Type Specific Gene Expression and Co-expression Network Architectures. Sci. Rep. 8, (2018).

24. Keren-Shaul, H. et al. A Unique Microglia Type Associated with Restricting Development of Alzheimer’s Disease. Cell 169, 1276–1290.e17 (2017).

25. Kosoy, R. et al. Alzheimer’s disease transcriptional landscape in ex vivo human microglia. Nat. Neurosci. 28, 1830–1843 (2025).

26. Lu, A. et al. Human microglial transitions at the Aβ–tau inflection point associate with divergent pathways to dementia and resilience. Nat. Med. (2026) doi:10.1038/s41591-026-04393-8.

27. Olah, M. et al. Single cell RNA sequencing of human microglia uncovers a subset associated with Alzheimer’s disease. Nat. Commun. 11, (2020).

28. Marschallinger, J. et al. Lipid-droplet-accumulating microglia represent a dysfunctional and proinflammatory state in the aging brain. Nat. Neurosci. 23, 194–208 (2020).

29. Haney, M. S. et al. APOE4/4 is linked to damaging lipid droplets in Alzheimer’s disease microglia. Nature 628, 154–161 (2024).

30. Baik, S. H. et al. A Breakdown in Metabolic Reprogramming Causes Microglia Dysfunction in Alzheimer’s Disease. Cell Metab. 30, 493–507.e6 (2019).

31. Gerrits, E. et al. Distinct amyloid-β and tau-associated microglia profiles in Alzheimer’s disease. Acta Neuropathol. 141, 681–696 (2021).

32. Xu, Z. et al. Microglia-specific regulation of lipid metabolism in Alzheimer’s disease revealed by microglial depletion in 5xFAD Mice. Nature Communications 16, (2025).

33. Claes, C. et al. Plaque-associated human microglia accumulate lipid droplets in a chimeric model of Alzheimer’s disease. Mol. Neurodegener. 16, (2021).

34. Patterson, M. T. et al. Trem2 promotes foamy macrophage lipid uptake and survival in atherosclerosis. Nature Cardiovascular Research 2, 1015–1031 (2023).

35. Gao, Q. et al. Role of iron in brain development, aging, and neurodegenerative diseases. Annals of Medicine vol. 57 Preprint at 10.1080/07853890.2025.2472871 (2025).

36. Ayton, S. et al. Brain iron is associated with accelerated cognitive decline in people with Alzheimer pathology. Mol. Psychiatry 25, 2932–2941 (2020).

37. Kenkhuis, B. et al. Iron accumulation induces oxidative stress, while depressing inflammatory polarization in human iPSC-derived microglia. Stem Cell Reports 17, 1351–1365 (2022).

38. Kenkhuis, B. et al. Iron loading is a prominent feature of activated microglia in Alzheimer’s disease patients. Acta Neuropathol. Commun. 9, (2021).

39. Van Duijn, S. et al. Cortical Iron Reflects Severity of Alzheimer’s Disease. Journal of Alzheimer’s Disease 60, 1533–1545 (2017).

40. Marschallinger, J. et al. Lipid-droplet-accumulating microglia represent a dysfunctional and proinflammatory state in the aging brain. Nat. Neurosci. 23, 194–208 (2020).

41. Kapralov, A. A. et al. Redox lipid reprogramming commands susceptibility of macrophages and microglia to ferroptotic death. Nat. Chem. Biol. 16, 278–290 (2020).

42. Ryan, S. K. et al. Microglia ferroptosis is regulated by SEC24B and contributes to neurodegeneration. Nat. Neurosci. 26, 12–26 (2023).

43. Marchand, B., Arsenault, D., Raymond-Fleury, A., Boisvert, F. M. & Boucher, M. J. Glycogen synthase kinase-3 (GSK3) inhibition induces prosurvival autophagic signals in human pancreatic cancer cells. Journal of Biological Chemistry 290, 5592–5605 (2015).

44. Parr, C. et al. Glycogen Synthase Kinase 3 Inhibition Promotes Lysosomal Biogenesis and Autophagic Degradation of the Amyloid-β Precursor Protein. Mol. Cell. Biol. 32, 4410–4418 (2012).

45. Beurel, E., Grieco, S. F. & Jope, R. S. Glycogen synthase kinase-3 (GSK3): Regulation, actions, and diseases. Pharmacology and Therapeutics vol. 148 114–131 Preprint at 10.1016/j.pharmthera.2014.11.016 (2015).

46. Huang, J., Guo, X., Li, W. & Zhang, H. Activation of Wnt/β-catenin signalling via GSK3 inhibitors direct differentiation of human adipose stem cells into functional hepatocytes. Sci. Rep. 7, (2017).

47. Avrahami, L. et al. Inhibition of glycogen synthase kinase-3 ameliorates β-amyloid pathology and restores lysosomal acidification and mammalian target of rapamycin activity in the alzheimer disease mouse model: In vivo and in vitro studies. Journal of Biological Chemistry 288, 1295–1306 (2013).

48. Licht-Murava, A., et al. P H A R M A C O L O G Y A Unique Type of GSK-3 Inhibitor Brings New Opportunities to the Clinic. https://www.science.org (2016).

49. Green, H. F. & Nolan, Y. M. GSK-3 mediates the release of IL-1β, TNF-α and IL-10 from cortical glia. Neurochem. Int. 61, 666–671 (2012).

50. Ríos, I. et al. Reprogramming of GM-CSF-dependent alveolar macrophages through GSK3 activity modulation. Elife 14, (2025).

51. Yousef, M. H., Salama, M., El-Fawal, H. A. N. & Abdelnaser, A. Selective GSK3β Inhibition Mediates an Nrf2-Independent Anti-inflammatory Microglial Response. Mol. Neurobiol. 59, 5591–5611 (2022).

52. Yuskaitis, C. J. & Jope, R. S. Glycogen synthase kinase-3 regulates microglial migration, inflammation, and inflammation-induced neurotoxicity. Cell. Signal. 21, 264–273 (2009).

53. Polychronopoulos, P. et al. Structural Basis for the Synthesis of Indirubins as Potent and Selective Inhibitors of Glycogen Synthase Kinase-3 and Cyclin-Dependent Kinases. J. Med. Chem. 47, 935–946 (2004).

54. Serenó, L. et al. A novel GSK-3β inhibitor reduces Alzheimer’s pathology and rescues neuronal loss in vivo. Neurobiol. Dis. 35, 359–367 (2009).

55. Bhujbal, S. P. et al. Gaining Insights into Key Structural Hotspots within the Allosteric Binding Pockets of Protein Kinases. International Journal of Molecular Sciences vol. 25 Preprint at 10.3390/ijms25094725 (2024).

56. Braak, H. & Braak, E. Acta H’ Pathologica Neuropathological Stageing of Alzheimer-Related Changes. Acta Neuropathol vol. 82 (1991).

57. Gabitto, M. I. et al. Integrated multimodal cell atlas of Alzheimer’s disease. Nat. Neurosci. 27, 2366–2383 (2024).

58. Morabito, S. et al. Single-nucleus chromatin accessibility and transcriptomic characterization of Alzheimer’s disease. Nat. Genet. 53, 1143–1155 (2021).

59. Lau, S.-F., Cao, H., Fu, A. K. Y. & Ip, N. Y. Single-nucleus transcriptome analysis reveals dysregulation of angiogenic endothelial cells and neuroprotective glia in Alzheimer’s disease. Proceedings of the National Academy of Sciences 117, 25800–25809 (2020).

60. Sadick, J. S. et al. Astrocytes and oligodendrocytes undergo subtype-specific transcriptional changes in Alzheimer’s disease. Neuron 110, 1788–1805.e10 (2022).

61. Zheng, G. X. Y. et al. Massively parallel digital transcriptional profiling of single cells. Nat. Commun. 8, (2017).

62. Hao, Y. et al. Integrated analysis of multimodal single-cell data. Cell 184, 3573–3587.e29 (2021).

63. Börner, K. et al. Anatomical structures, cell types and biomarkers of the Human Reference Atlas. Nature Cell Biology vol. 23 1117–1128 Preprint at 10.1038/s41556-021-00788-6 (2021).

64. Wolf, F. A., Angerer, P. & Theis, F. J. SCANPY: Large-scale single-cell gene expression data analysis. Genome Biol. 19, (2018).

65. de Leeuw, C. A., Mooij, J. M., Heskes, T. & Posthuma, D. MAGMA: Generalized Gene-Set Analysis of GWAS Data. PLoS Comput. Biol. 11, (2015).

66. Liberzon, A. et al. The Molecular Signatures Database Hallmark Gene Set Collection. Cell Syst. 1, 417–425 (2015).

67. Robinson, M. D., McCarthy, D. J. & Smyth, G. K. edgeR: A Bioconductor package for differential expression analysis of digital gene expression data. Bioinformatics 26, 139–140 (2009).

68. Bergen, V., Lange, M., Peidli, S., Wolf, F. A. & Theis, F. J. Generalizing RNA velocity to transient cell states through dynamical modeling. Nat. Biotechnol. 38, 1408–1414 (2020).

69. Wong, K. G. et al. CryoPause: A New Method to Immediately Initiate Experiments after Cryopreservation of Pluripotent Stem Cells. Stem Cell Reports 9, 355–365 (2017).

70. Haenseler, W. et al. A Highly Efficient Human Pluripotent Stem Cell Microglia Model Displays a Neuronal-Co-culture-Specific Expression Profile and Inflammatory Response. Stem Cell Reports 8, 1727–1742 (2017).

71. Delignette-Muller, M. L., Siberchicot, A., Larras, F. & Billoir, E. M E R S E N N E Peer Community Journal Section: Ecotoxicology and Environmental Chemistry DRomics, a workflow to exploit dose-response omics data in ecotoxicology. (2023) doi:10.24072/pci.

72. ICell ® Microglia Application Protocol Labeling Amyloid Beta with PHrodo Red. https://fujifilmcdi.com/assets/CDI_iCellMGL_Incucyte_AP.pdf.

