## Supplementary Figures for "Characterization and pharmacological modulation of Alzheimer’s disease-associated human microglial states"

### Extended Data Figure 1

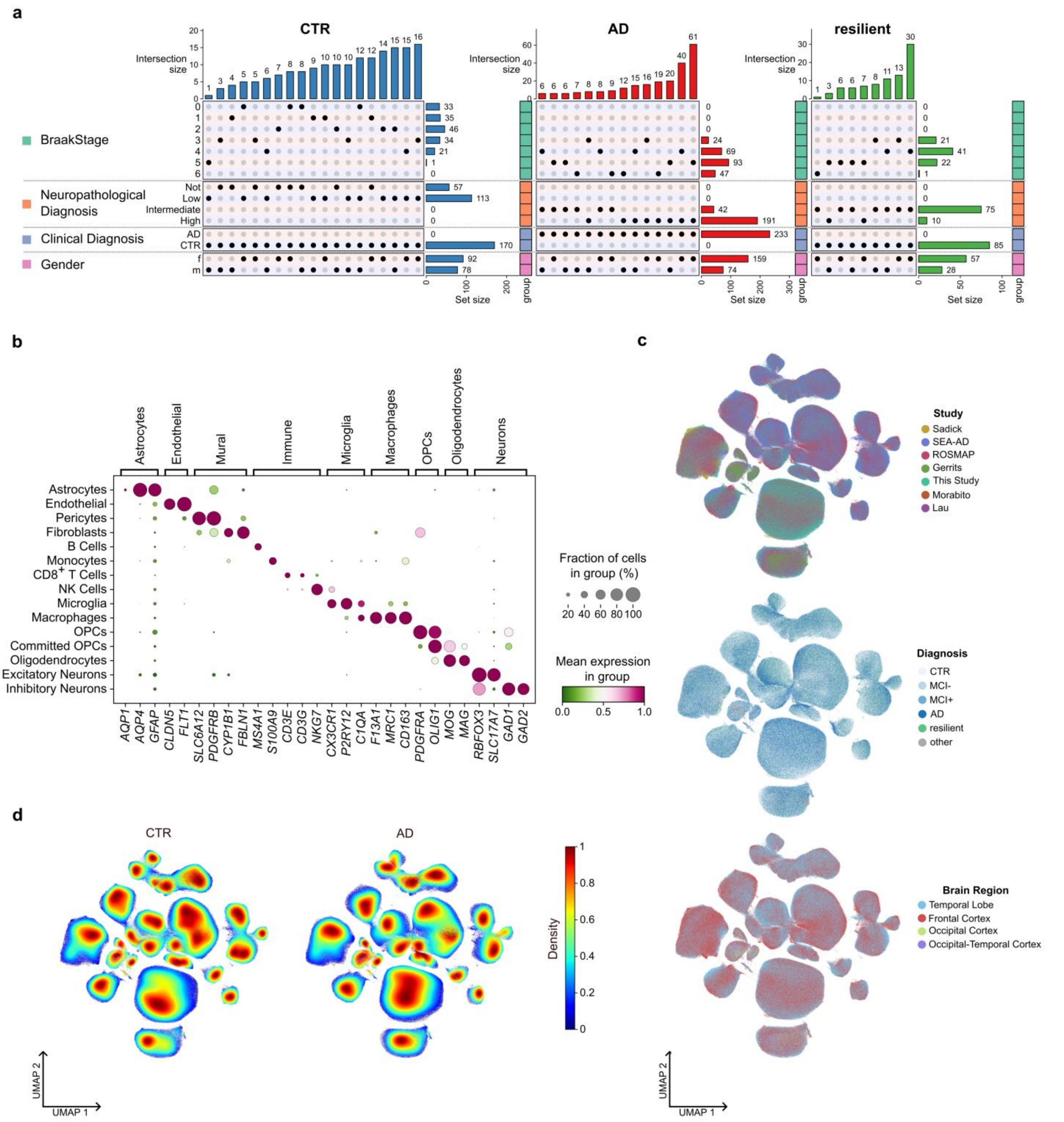


**Extended Data Fig. 1 | Alzheimer's disease single-nucleus transcriptomic atlas cohort composition and data integration**

**a)** UpSet plots summarizing patient metadata combinations within each diagnostic group (CTR, AD and resilient). Dots indicate membership in each metadata category (Braak stage, neuropathological diagnosis, clinical diagnosis and gender), vertical bars indicate intersection size and horizontal bars indicate set size. **b)** Dot plot of key marker genes used to support major cell type annotation across the atlas. Dot size indicates the fraction of cells expressing each gene within a cell type and color indicates mean expression level. **c)** UMAPs of all nuclei from the integrated snRNA-seq atlas (same as in Fig. 1a) colored by data source (left), joint diagnosis (middle), and tissue brain region of origin (right). **d)** Density visualization showing the distribution of nuclei across the UMAP embedding separated by clinical diagnosis (top: CTR and bottom: AD).

### Extended Data Figure 2

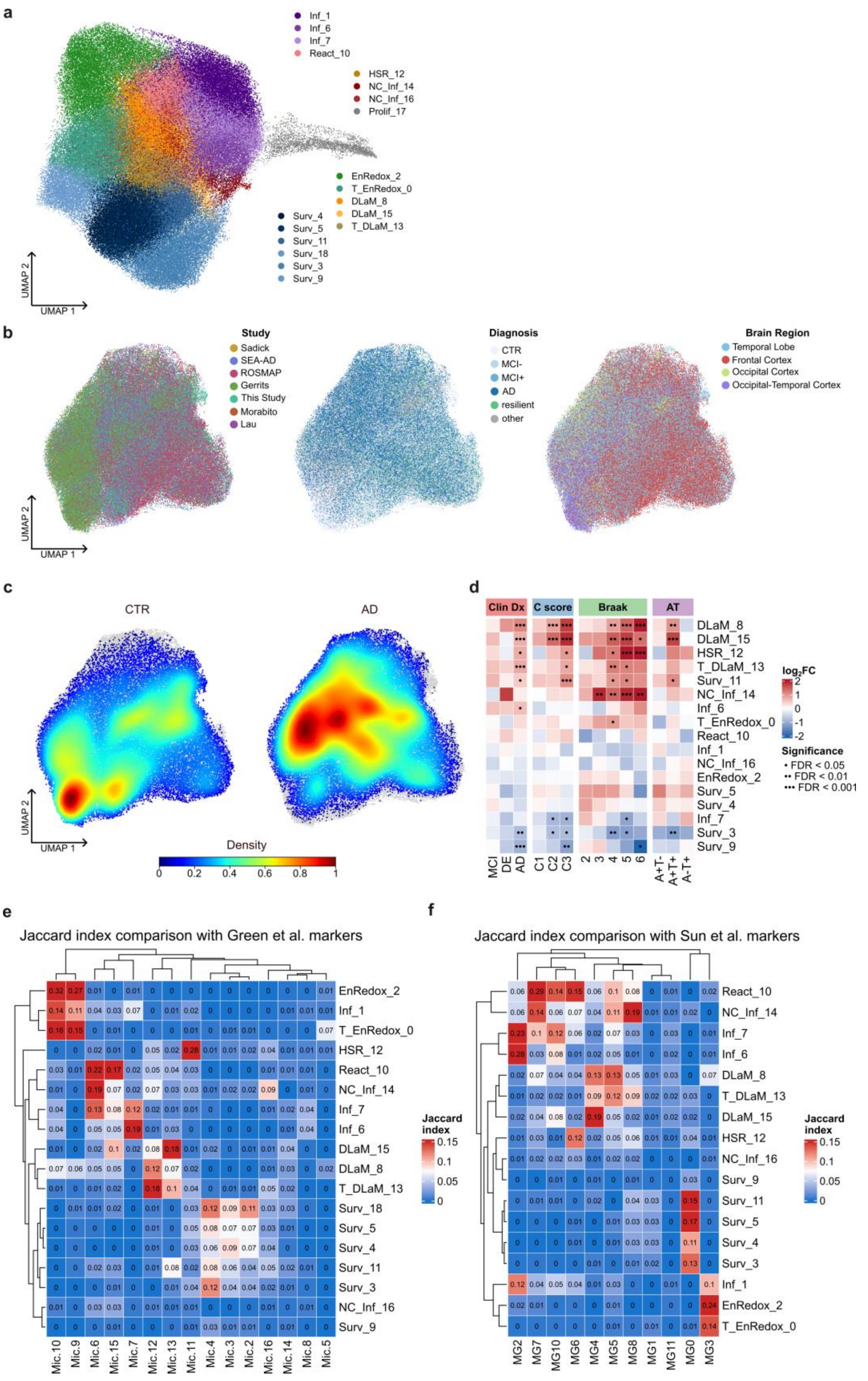


**Extended Data Fig. 2 | Microglia object data structure and comparison to published microglia states.**

**a)** UMAP embedding of reclustered microglia object from Fig. 2a including the proliferative cluster 17. **b)** UMAPs of microglia as in Fig. 2a colored by data source (left), joint diagnosis (middle), and tissue brain region of origin (right). **c)** Density visualization showing the distribution of nuclei across the UMAP embedding separated by clinical diagnosis (top: CTR and bottom: AD). **d)** Heatmap summarizing associations between microglial states and clinical/neuropathological variables. Color denotes the direction and magnitude of the association. (empirical Bayes quasi-likelihood F-tests corrected for confounders, (·) = FDR < 0.05, (··) = FDR < 0.01, (···) = FDR < 0.001). **e-f)** Heatmaps of Jaccard indices for comparisons between microglia states identified in this study with those reported by Green et al. (d) and Sun et al. (e). See section "Microglia cluster similarities across studies" in Methods for further details.

### Extended Data Figure 3

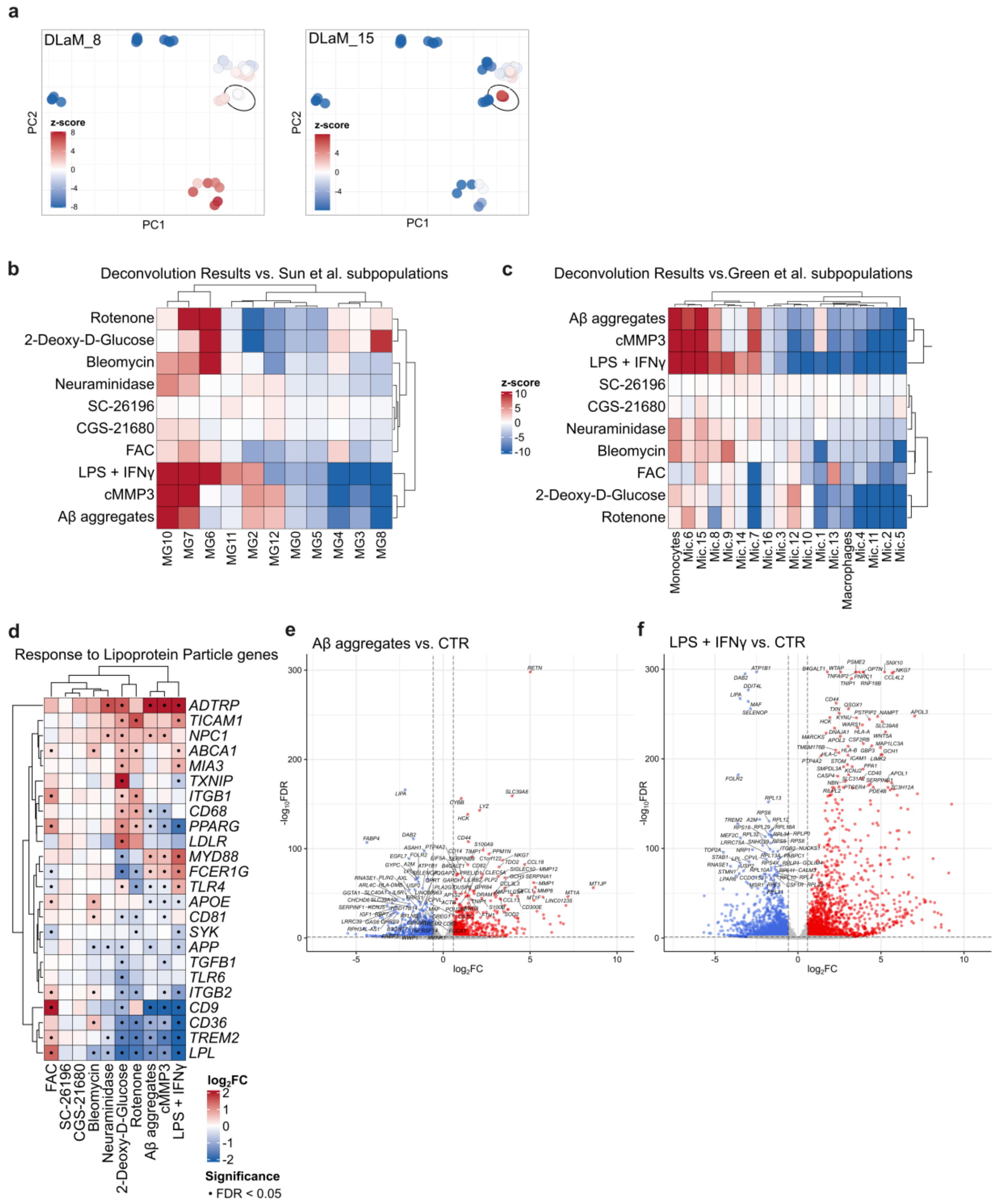


**Extended Data Fig. 3 | Further characterization of stimulated hiPSC microglia transcriptional responses**

**a)** PCA of stimulation-induced transcriptional changes (same as in Fig. 3b) colored by DLaM cluster similarity (z-score), shown separately for DLaM_15 (top) and DLaM_8 (bottom). Highlighted points indicate FAC stimulation. **b-c)** Deconvolution to patient microglia states as reported by Green et al. (a) and Sun et al. (b) parallel to Fig. 3c. Heatmaps show BRETIGEA-based normalized z-scores (see Methods) using microglia cluster markers from the respective studies. **d)** Expression heatmap of genes belonging to the GO term "Response to Lipoprotein Particle" (GO:0055094) across stimulation conditions from Fig. 3. Statistical test as determined by limma-voom pipeline, see Methods, (·) = FDR < 0.05). **e-f)** Aβ aggregates (d) and LPS+IFNγ (e) induced transcriptional signatures shown as Volcano plots of the respective stimuli vs. solvent control (as determined by limma-voom pipeline, see Methods).

### Extended Data Figure 4

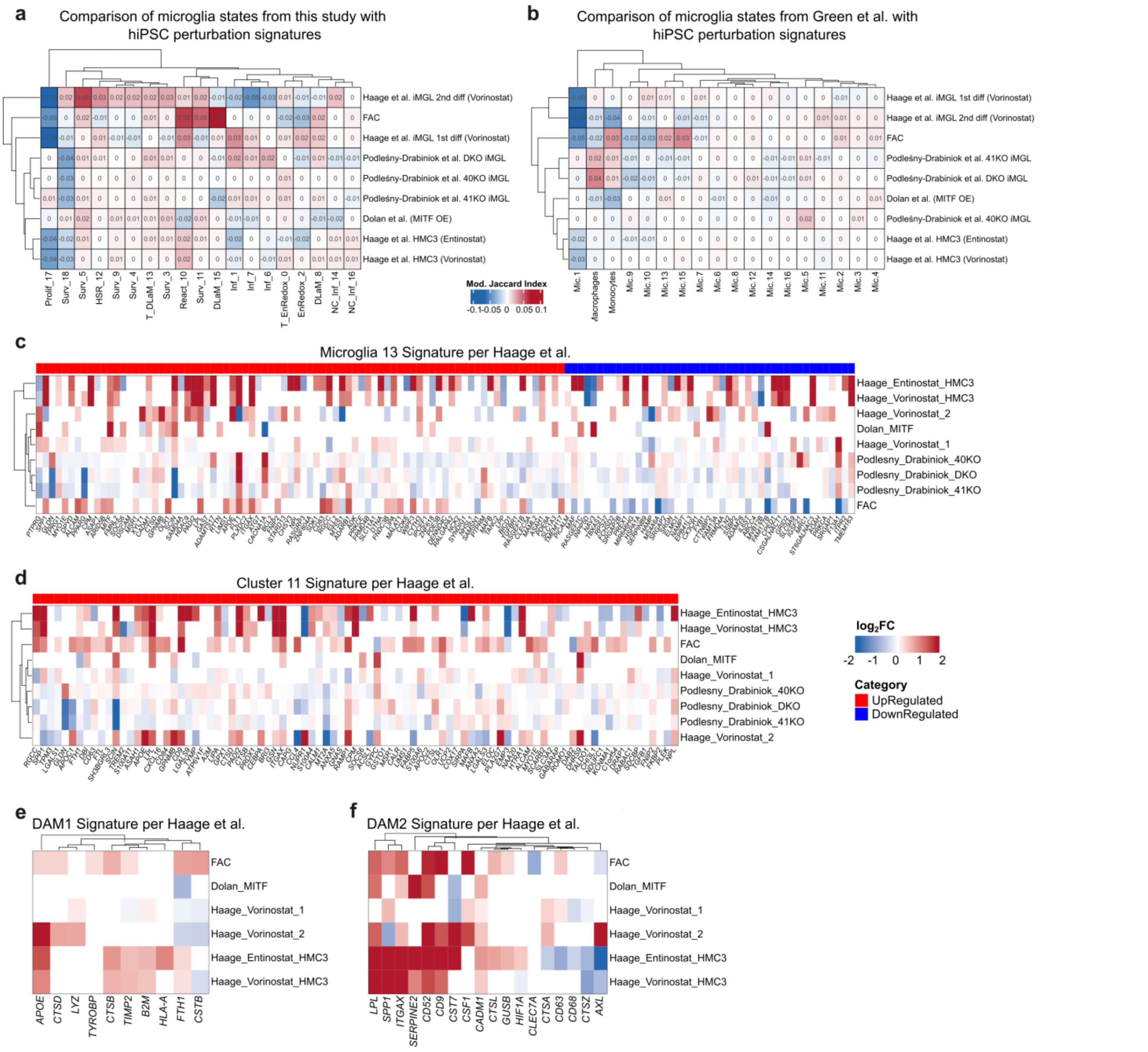


**Extended Data Fig. 4 | Comparison of FAC-induced transcriptional signature to published hiPSC microglia perturbation signatures**

**a-b)** Heatmaps with modified Jaccard indices highlight overlaps between patient microglia state marker genes from this study (a) and Green et al. (b) with different hiPSC microglia perturbation signatures from this study (FAC) and different publications[7-9]. See section "Microglia cluster and stimulation signature similarities" in Methods for further details. **c-f)** Heatmaps of published patient disease associated microglia states (source indicated above each heatmap) across different perturbations, FAC (this study) and published data as indicated. Color indicates signed log2FC (scale shown next to heatmap in (d) common for all four). For (c) and (d), the colored top bar annotation shows whether genes are upregulated (red) or downregulated (blue) within the respective patient microglia state.

#### Extended Data Figure 5

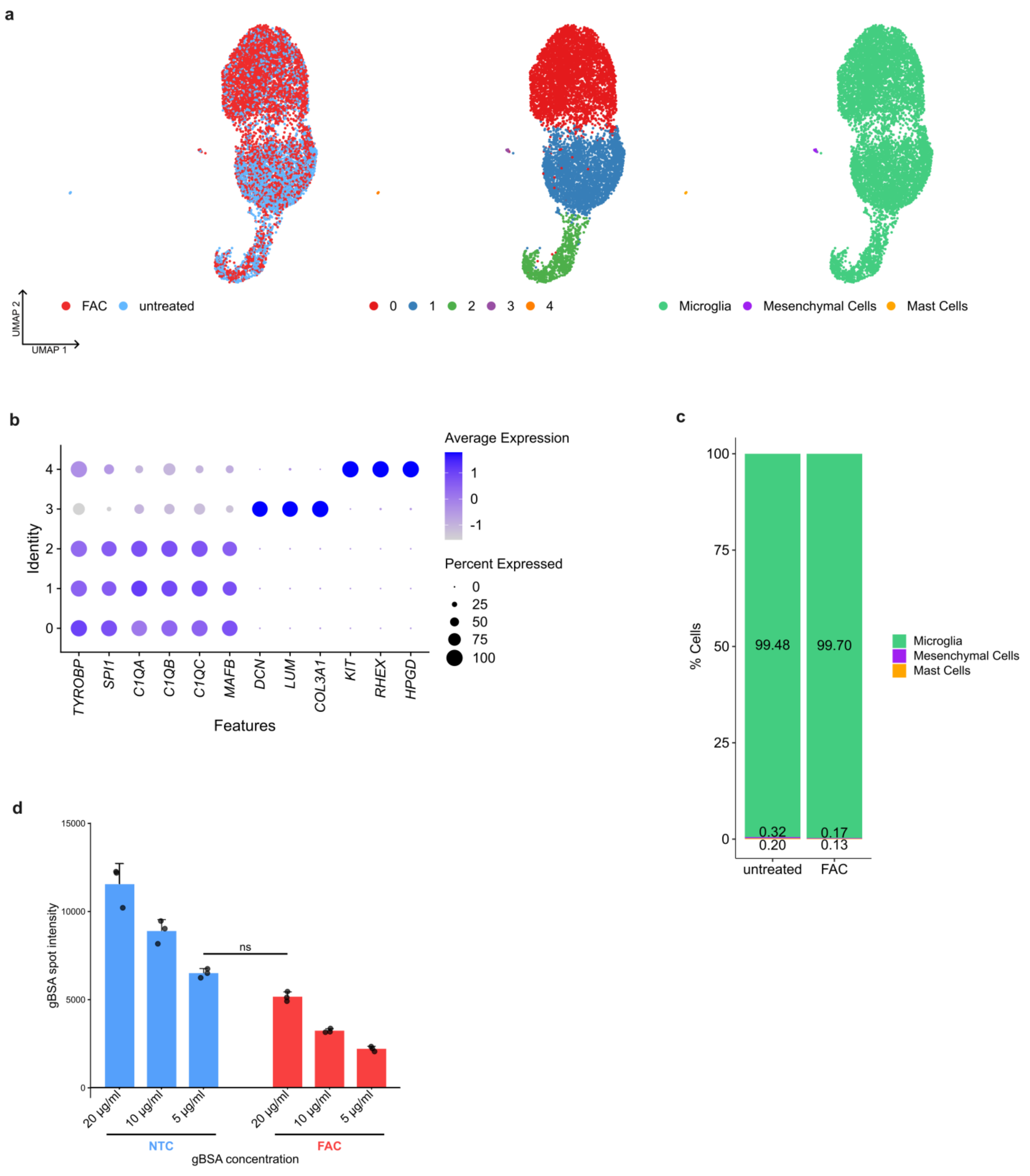


**Extended Data Fig. 5 | Quality control of hiPSC microglia culture scRNA-seq data and titration of BSA uptake**

**a)** UMAPs of scRNA-seq data from untreated and FAC-treated hiPSC microglia cultures with mesenchymal and mast cells included. Colored by treatment (left), clusters (middle), and major cell type annotation (right). **b)** Dot plot of cluster marker gene expression for the hiPSC microglia culture clusters shown in (a). **c)** Major cell type composition of untreated and FAC-treated hiPSC microglia cultures as identified in the scRNA-seq data showing the fraction of microglia, mesenchymal cells, and mast cells per condition. **d)** Quantification of gBSA uptake at the start of the pulse-chase assay across gBSA concentrations in untreated (NTC, blue) and FAC-treated (red) cultures, used to match uptake between conditions for the lysosomal activity measurements sown in Fig. 4f. No statistically significant difference in gBSA uptake at 5 µg/ml in unstimulated compared to 20 µg/ml in FAC-stimulated hiPSC microglia using a two-way ANOVA with follow-up Tukey's multiple comparison test (main gBSA-Concentration effect $F[2,12] = 74.37$, $p < 0.0001$, $\eta 2p = 0.256$; main Treatment effect $F[1,12] = 409.8$, $p < 0.0001$, $\eta 2p = 0.705$ and interaction Concentration × Treatment $F[2,12] = 5.19$, $p < 0.0001$, $\eta 2p = 0.017$. Data are presented as mean ± SD with n = 3.

#### Extended Data Figure 6

a

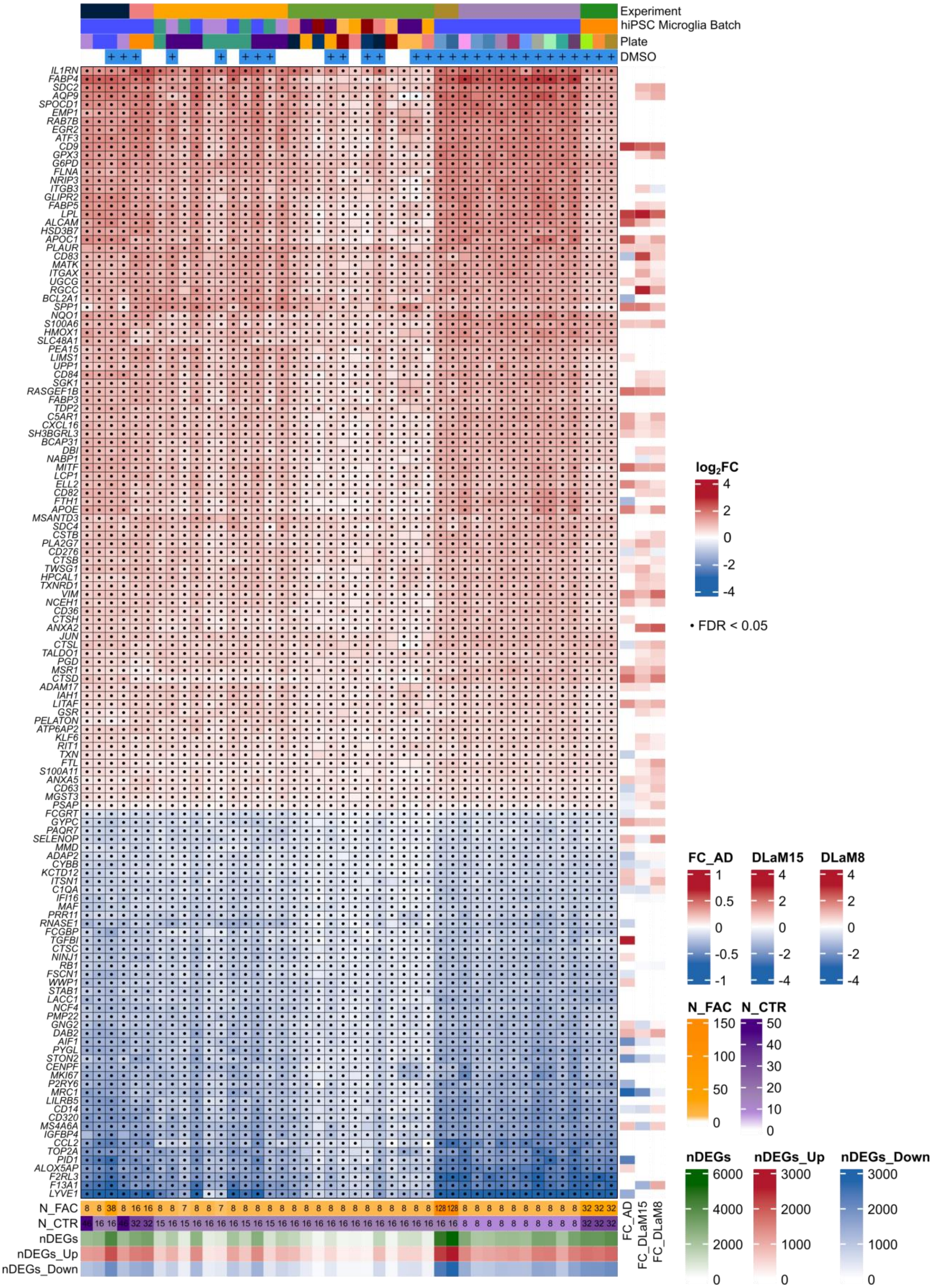


**Extended Data Fig. 6 | Robustness and reproducibility of the FAC-induced transcriptomic signature**

**a)** Heatmap of the most consistently FAC-induced DEGs across seven independent experiments. Top annotation bars contain sample descriptors, i.e., experiment, cell batch, plate, and DMSO treatment status for each column. Right-side annotation shows log2FC for respective genes in patient microglia, separately for AD vs. healthy controls, DLaM15 cluster vs. all other patient microglia clusters, and DLaM8 cluster vs. all other patient microglia clusters. Bottom annotation bars report sample numbers (N_FAC and N_CTR) as well as numbers of all DEGs (green), upregulated DEGs (red) and downregulated DEGs (blue) for each FAC vs. CTR comparison. A (·) indicates genes that are significantly changed compared to unstimulated controls with FDR < 0.05, as determined by limma-voom, FDR < 0.05, no log2FC filtration.

### Extended Data Figure 7

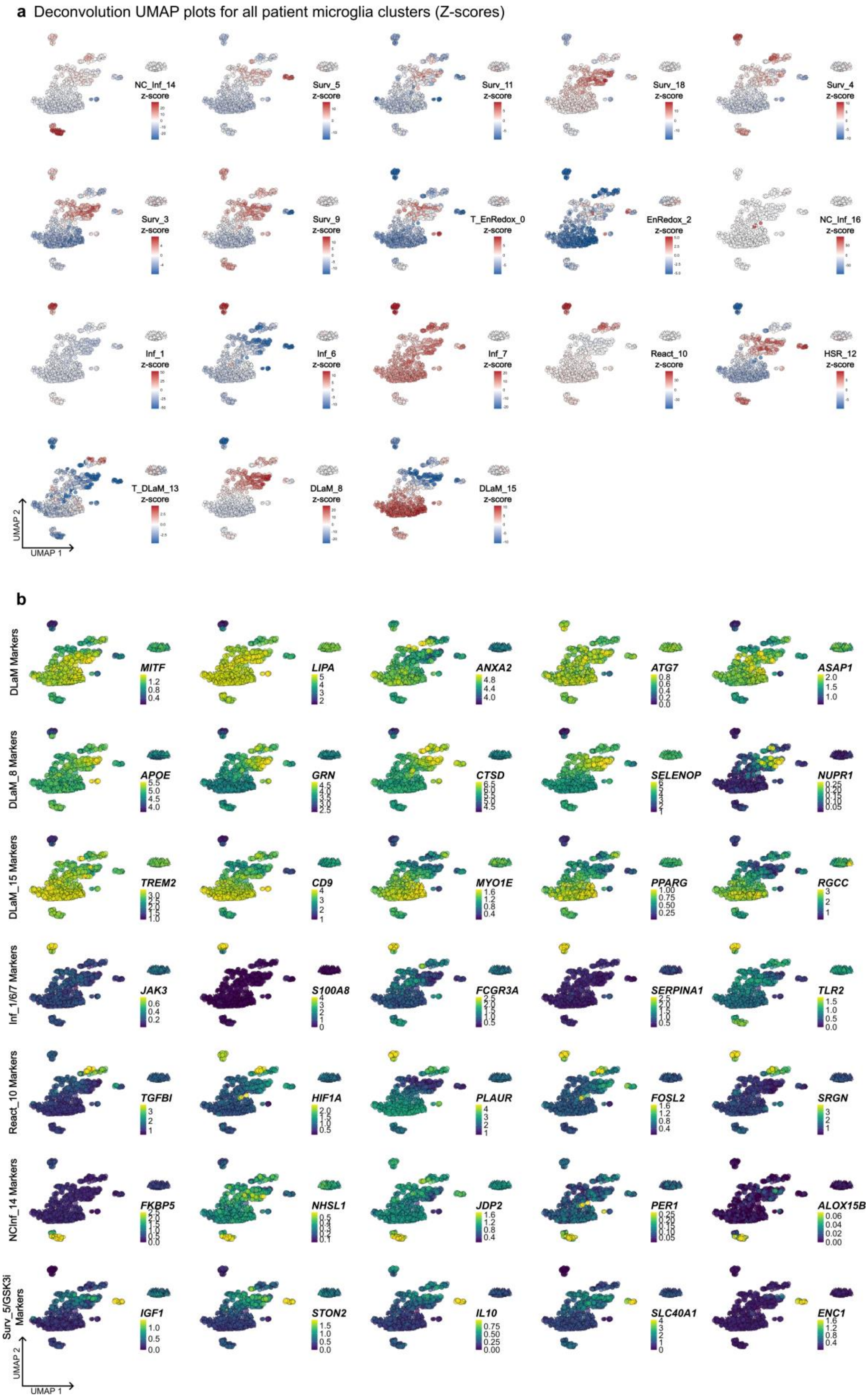


**Extended Data Fig. 7 │ Deconvolution of compound perturbations to patient microglia states**

**a)** UMAPs (same UMAP as in Fig. 5b) overlayed with deconvolution scores for patient microglia clusters identified in this study. **b)** UMAPs overlayed with expression of representative marker genes from patient microglia clusters identified in this study, as well as selectively up-regulated genes by GSK3 inhibitors.

### Extended Data Figure 8

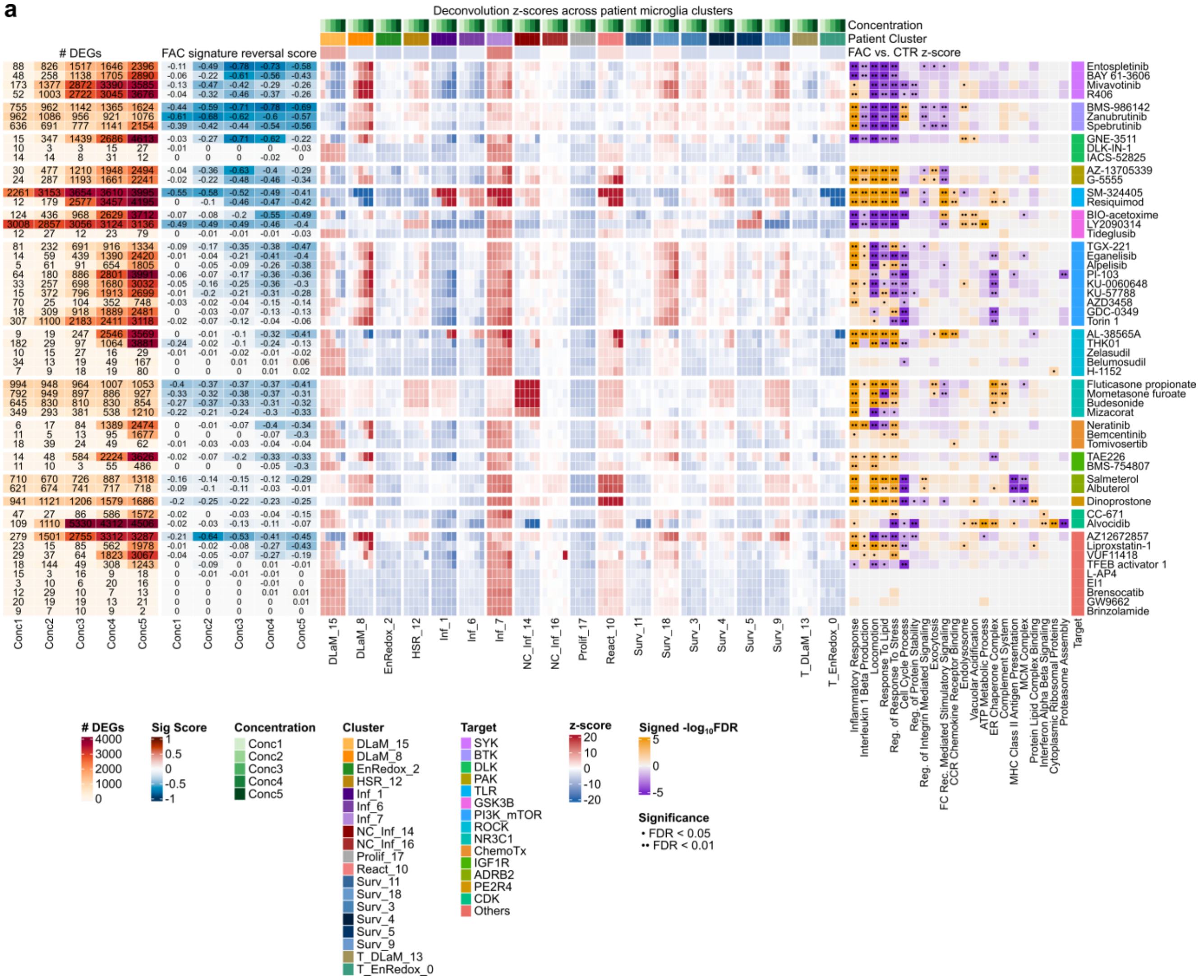

**Extended Data Fig. 8 │ Detailed characterization of tanscriptomics-based small molecule screening**

**a)** Heatmaps summarize number of DEGs (as determined by limma-voom, FDR < 0.05, no log2FC filtration), FAC signature-reversal score (σ), averaged deconvolution scores, and signed –log10FDR estimates from pathway enrichment analysis across all compounds. Pathway enrichment was performed using a hypergeometric test, including both up- and down-regulated DEGs, and applying the Benjamini-Hochberg method for multiple hypothesis correction. Genes were annotated as up-regulated by a compound if significantly increasing in at least 2 concentrations or down-regulated by a compound if significantly decreasing in at least 2 concentrations (as determined by limma-voom pipeline FDR < 0.05, see Methods). A (·) indicates pathways that are significantly enriched by a given compound with FDR < 0.05. The color of each cell in the heatmap was determined by the sign of the median of the average per-gene log2FC across all DEGs in each pathway. Accompanying annotation bars indicate compound concentrations, patient microglia clusters, and averaged FAC deconvolution z-scores by patient microglia cluster (all top), and annotated compound targets (right).

### Extended Data Figure 9

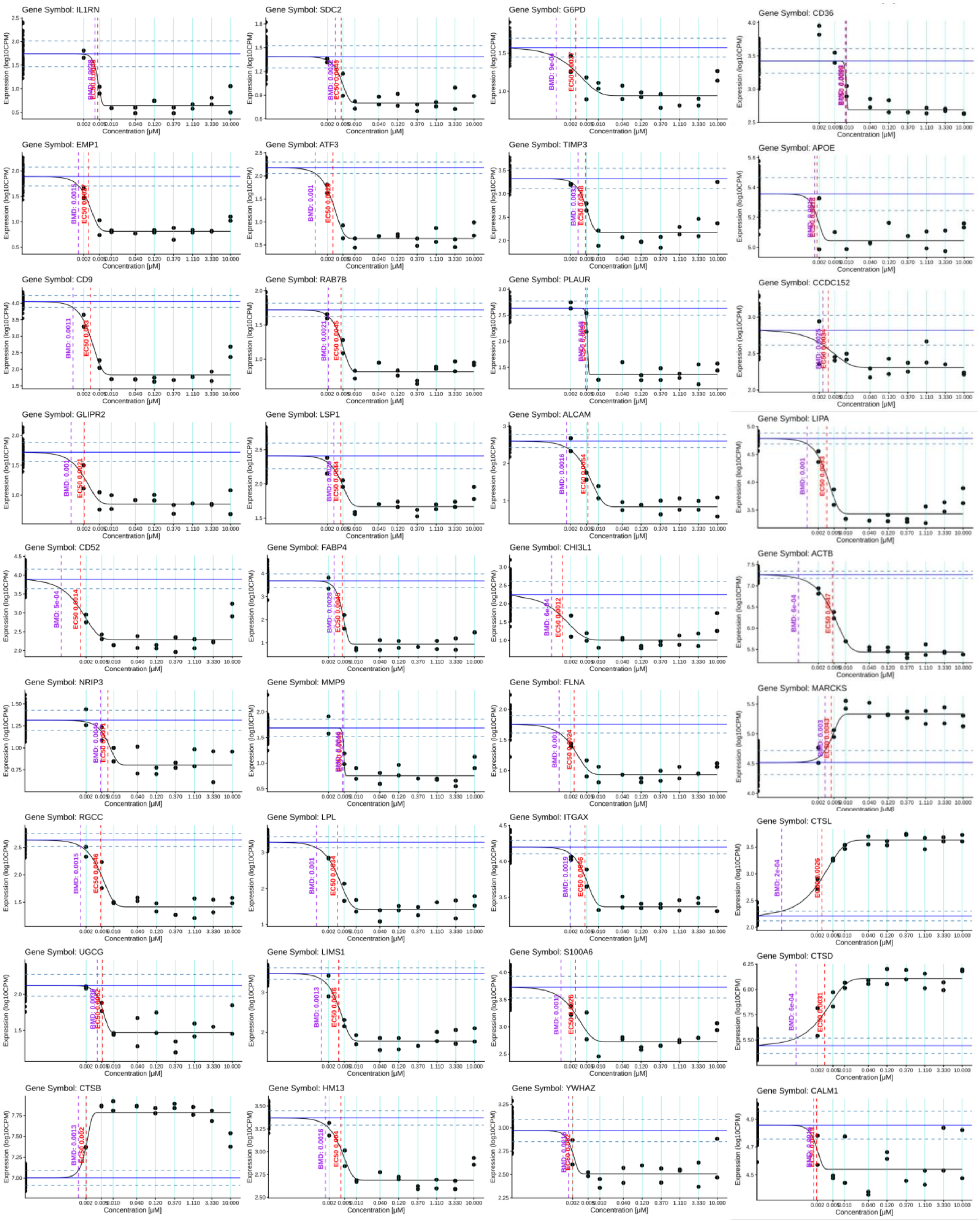


**Extended Data Fig. 9 | Extended example concentration response curves for genes regulated by LY2090314**

Additional example concentration response curves for genes regulated by treatment with LY2090314 as in Fig. 7a. (see Methods)

### Extended Data Figure 10

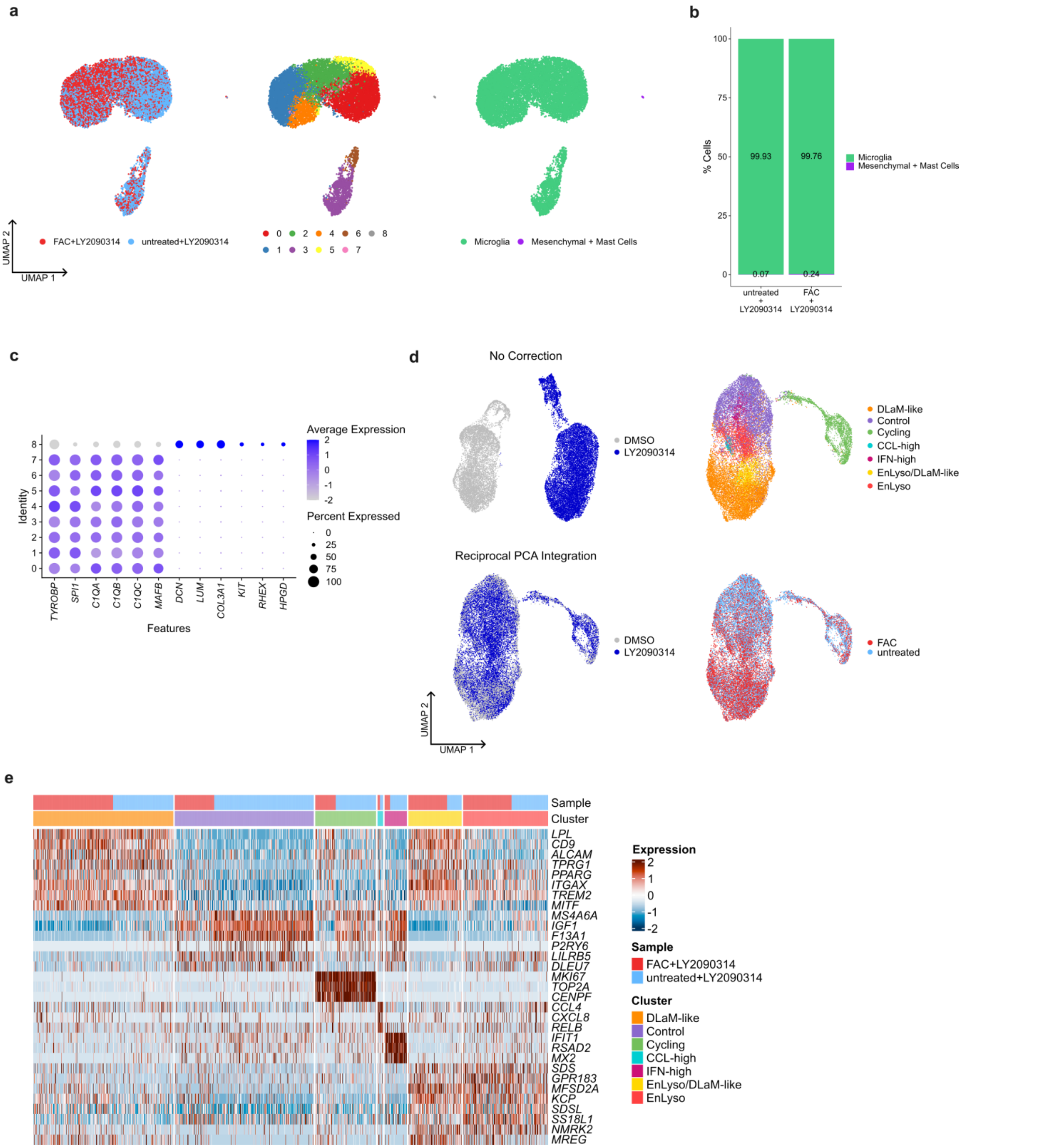


**Extended Data Fig. 10 | Quality control of hiPSC microglia with LY2090314 treatment scRNA-seq data**

**a)** UMAPs of scRNA-seq data from untreated and FAC-treated hiPSC microglia cultures with LY2090314 treatment with mesenchymal and mast cells included. Colored by treatment (left), clusters (middle), and major cell type annotation (right). **b)** Major cell type composition of untreated and FAC-treated hiPSC microglia cultures with LY2090314 treatment as identified in the scRNA-seq data showing the fraction of microglia, mesenchymal and mast cells per condition. **c)** Dot plot of cluster marker gene expression for the hiPSC microglia culture clusters shown in (a) and (b). **d)** UMAPs of scRNA-seq data from untreated and FAC-treated hiPSC microglia with and without LY2090314 treatment. Integration without sample correction and colored by LY2090314 treatment is shown in upper left. Integration with reciprocal PCA integration colored by LY2090314 treatment (bottom left), clusters (upper right) and treatment (bottom right). **e)** Heatmap of cluster markers and their expression in individual cells grouped by cluster and sample.
